# Homeostatic fibroblast niche collapse in clinically uninvolved skin defines a subclinical state of systemic sclerosis

**DOI:** 10.64898/2026.11.05.764714

**Authors:** Alex K. Hu, Lin Wei Tung, Thomas Pulliam, Cory L. Simpson, Sandra Lord, Daniel E. Furst, Jeffrey S. Carlin, Hannah A. DeBerg, Daniel J. Campbell, Peter A. Morawski

## Abstract

Systemic sclerosis, or scleroderma, is a chronic autoimmune disease characterized by vasculopathy, immune dysregulation, and progressive fibrosis of the skin and internal organs. Heterogeneous clinical presentation and polygenic risk complicate our understanding of disease mechanisms and the path to effective therapy. Single-cell transcriptomic analyses have revealed substantial heterogeneity in skin, but disease-associated reprogramming remains poorly defined. To address this, we performed high-resolution spatial transcriptomic profiling of paired clinically uninvolved and clinically affected scleroderma donor skin alongside healthy controls. We identified a continuum of cellular, molecular, and spatial changes to the stroma that were already evident in clinically uninvolved tissue yet remained distinct from the more extensive changes in paired fibrotic biopsies. Uninvolved skin was comparable to healthy tissue by clinical and histopathological assessment, but distinguishable by the proportional reduction, transcriptional reprogramming, and spatial isolation of LGR5^+^SCARA5^+^PI16^+^ progenitor fibroblasts. In parallel, the dermal vascular smooth muscle compartment shifted from a contractile to a synthetic, activated state, suggesting dysfunctional vascular reprogramming occurs in the absence of overt fibrosis. Cell neighborhood analysis revealed that progenitor fibroblasts occupy dense stromal niches in healthy skin that collapse in clinically uninvolved tissue, replaced by isolated, disease-associated fibroblast populations. These data indicate that clinically uninvolved skin occupies an intermediate subclinical state of scleroderma, establishing a framework for strategies to allow early diagnosis and preserve homeostatic fibroblast niches to prevent irreversible tissue remodeling and fibrotic commitment.

**One sentence summary:** Clinically uninvolved scleroderma skin reveals a subclinical spatial-molecular state defined by progenitor fibroblast niche collapse.

## INTRODUCTION

Systemic sclerosis (SSc), also called scleroderma, is an idiopathic autoimmune disease characterized by vasculopathy, dysregulated immune cell function, and connective tissue fibrosis. It has no cure and among the highest mortality rates of any rheumatic disease. SSc is polygenic and highly heterogeneous in its clinical presentation, which complicate diagnosis, prognosis, therapeutic algorithms, and drug design (*1–3*). Two major disease subsets, limited cutaneous systemic sclerosis (lcSSc) and diffuse cutaneous systemic sclerosis (dcSSc), are primarily distinguished by the anatomical regions of skin impacted and associated serum autoantibody profiles (*4*). Although dcSSc generally demonstrates more extensive fibrosis and higher overall mortality, individuals with lcSSc frequently develop severe systemic complications including pulmonary arterial hypertension, vasculopathy, digital ischemia and ulceration, and gastrointestinal dysmotility, resulting in substantial morbidity and mortality (*5*). Skin involvement, measured using the modified Rodnan skin score (mRSS), correlates with systemic disease activity early in dcSSc but otherwise has limited prognostic value (*6–8*). Despite representing approximately 60-70% of scleroderma cases, lcSSc patients are excluded from most skin-targeting clinical trials (*9, 10*), a central gap in patient care that underscores a need for improved understanding of disease states. Sampling adjacent uninvolved and affected skin in SSc would provide a unique opportunity to evaluate the mechanisms underlying the development of fibrosis.

Vasculopathy and inflammatory immune cell activity are among the early manifestations of SSc and precede commitment to fibrosis, yet the dermal vasculature and its surrounding contractile smooth muscle remain poorly understood (*11–13*). Dermal fibroblasts by comparison have been extensively characterized owing to their abundance and central role in fibrosis. Single-cell transcriptomic analyses have shown that fibroblasts are compositionally diverse at steady-state and dynamically reprogrammed during SSc (*14–18*). LGR5^+^ progenitor fibroblasts enriched in healthy skin are progressively lost in SSc and replaced by pathogenic myofibroblasts marked by pro-fibrotic genes such as COMP, POSTN, and SFRP4 (*14, 16, 17, 19, 20*). LGR5, a Wnt-pathway stem cell marker, is co-expressed with SCARA5 and PI16 on a subset of the conserved ‘universal’ fibroblast lineage, which is implicated as the precursor pool for cells involved in tissue homeostasis, extracellular matrix maintenance, and wound healing (*19–23*). Despite evidence of vascular, immune, and stromal involvement in SSc, patient care remains hindered by an incomplete understanding of the spatial-molecular changes that initiate and advance pathology (*24*). Determining if lesions are stable or if nearby normal skin remains at risk could inform the dose and duration of high-risk immunosuppressive therapies.

Trajectory modeling of fibroblast transcriptional states suggests that POSTN^+^PRSS23^+^COMP^+^ myofibroblasts arise from a SCARA5^+^CD34^+^ progenitor pool and adopt distinct spatial organization patterns in dcSSc relative to healthy skin (*16*). These inferences, however, rest on dissociated or low-resolution data, and none have examined clinically uninvolved skin. Paired affected and uninvolved skin samples from the same donor offers a unique opportunity to resolve transcriptional and spatial changes that emerge in a subclinical state, yet uninvolved skin has rarely been examined in scleroderma. The few studies to do so used bulk-tissue microarrays with limited resolution of cellular composition or spatial organization – a dcSSc study concluding uninvolved and lesional skin were indistinguishable (*25*); a study of clinically uninvolved forearm skin in established lcSSc that identified molecular subsets distinguished by extracellular matrix, TGF-β response, and cardiovascular development genes (*26*); and an analysis of morphea that pooled uninvolved skin with healthy controls (*27*). Thus, whether clinically susceptible but as-yet uninvolved skin represents an early or intermediate disease state has not been definitively addressed.

To examine the changing cell neighborhoods and transcriptional programs of scleroderma, we performed single-cell spatial transcriptomics on clinically affected lcSSc skin and proximal, but clinically uninvolved tissue from the same donors, compared with age- and sex-matched healthy controls. We show that clinically uninvolved tissue occupies an intermediate state during fibrotic disease, between affected and healthy skin, as defined by: (i) changes in progenitor fibroblast proportions, transcriptional programs, and spatial position; (ii) reprogramming of dermal vascular smooth muscle from a contractile toward a synthetic, activated state; and (iii) homeostatic fibroblast niche collapse and progressive isolation of remaining progenitor cells. We integrated these features into a spatial-molecular index that ordered samples along a clinical continuum, distinguishing clinically uninvolved skin from both fibrotic skin and healthy controls. Our data support a model in which homeostatic fibroblast niche collapse defines a subclinical state of lcSSc that occurs independently of pathogenic myofibroblast expansion and tissue fibrosis, supporting the potential utility of strategies aimed at reinforcing or restoring progenitor fibroblasts, alongside existing biologic and anti-fibrotic regimens.

## RESULTS

### Spatially resolved single-cell profiling maps the cellular landscape of paired uninvolved and affected lcSSc skin

To map the cellular composition and gene programs of lcSSc donor tissue across a range of clinical involvement, we performed high-resolution spatial transcriptomic profiling (Visium HD; 10X Genomics) of 24 skin biopsies obtained from 15 individuals: paired clinically uninvolved (SU) and clinically affected (SA) skin from 9 lcSSc donors alongside 6 age- and sex-matched healthy control (HC) samples (**Table S1, Fig. 1A**). Given the typical distal-to-proximal spread of lcSSc, HC and SU samples were taken from the forearm and SA samples from the finger in seven of nine donors. Affected and uninvolved skin regions were determined at the time of sample collection by a dermatologist and enabled within-donor comparisons of features that emerge in the presence or absence of overt pathology. Clinically uninvolved samples and site-matched healthy control forearm skin were indistinguishable by blinded histologic assessment (**Fig. 1B, S1A**), though a few from each group contained areas of mild fibrosis. Clinically affected samples exhibited moderate-to-severe fibrosis throughout the tissue, consistent with their mRSS (**Fig. S1B**).

**Figure 1.**
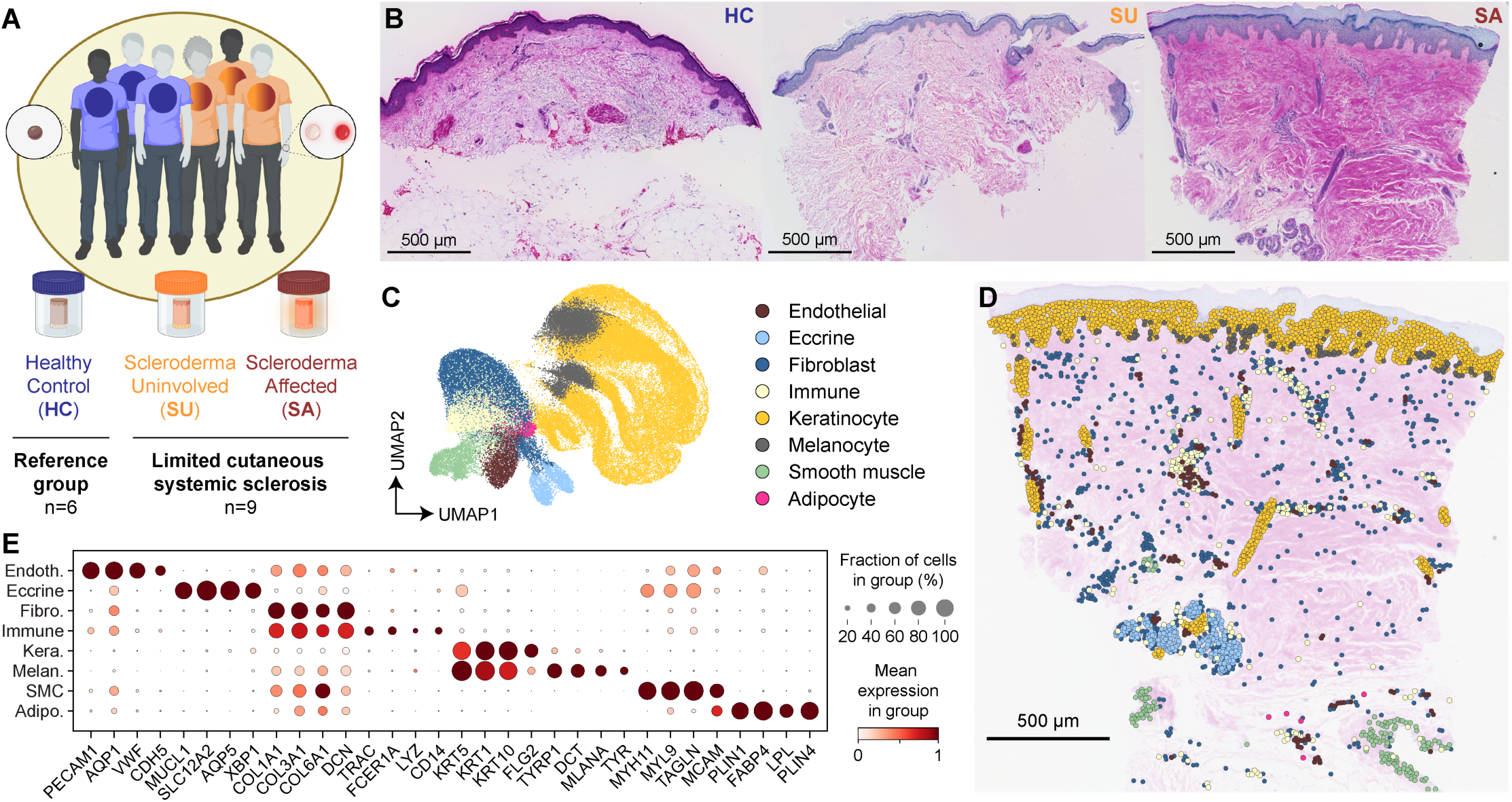
Single-cell spatial profiling of healthy, clinically uninvolved and affected lcSSc skin. (**A**) Study cohort: 24 skin punch biopsies from 15 donors (6 healthy controls, HC; 9 lcSSc donors contributing paired clinically uninvolved, SU, and affected, SA, skin), profiled by 10X Genomics Visium HD. (**B**) Representative H&E-stained biopsy sections per group. (**C**) UMAP of 96,865 segmented cells colored by eight distinct cutaneous populations. (**D**) Representative section with segmented cells overlaid on H&E image, colored as in (C), recovering epidermal, dermal, and hypodermal populations. (**E**) Dot plot of lineage markers (dot size, fraction of cells expressing; color, scaled mean expression). Scale bars, 500 µm.

We clustered 96,865 segmented cells from all samples using highly variable gene expression and visualized them by uniform manifold approximation and projection (UMAP), resolving eight populations: keratinocytes, fibroblasts, endothelial cells, immune cells, melanocytes, smooth muscle cells (SMC), adipocytes, and eccrine cells (**Fig. 1C**). Populations were annotated using both spatial position and lineage-defining gene expression (**Fig. 1D**, **1E**) and localized reproducibly to epidermal, dermal, and hypodermal skin regions, including dermal adnexal structures and pilosebaceous units. Cellular composition was variable between samples, but largely conserved across groups, save for a few differences between HC and SA, such as reduced fibroblast frequency (**Fig. S1C**). The immune compartment contained T cells and macrophages, while the endothelial compartment contained both vascular and lymphatic endothelial cells (VEC, LEC) (**Fig. S1D-S1F**). Thus, SU tissue showed no significant broad cell-type differences from site-matched HC skin.

### Fibroblast compositional shifts in clinically uninvolved skin define a subclinical state in lcSSc

We performed independent clustering of 27,000 dermal fibroblasts from all 24 samples, identifying nine transcriptionally distinct populations (Fb1-Fb9), present across study groups at varying frequencies (**Fig. 2A**, **2B**). Cell annotations were determined using differentially expressed genes per cluster: perivascular (Fb1; CXCL12, APOE, IGFBP7), inflammatory fibroblastic reticular cell (FRC) (Fb2; CD74, PTGDS), LGR5^+^SCARA5^+^PI16^+^ progenitor (Fb3), matrix-producing (Fb4; MMP2, COMP), superficial papillary (Fb5; APCDD1), fibrotic myofibroblast (Fb6; CTHRC1, POSTN), resting (Fb7; DCN), reticular (Fb8; SFRP1), and Schwann-like (Fb9; NGFR, MPZ) (**Fig. 2C**), while spatial mapping revealed ordered, reproducible dermal distribution (**Fig. 2D**). Clusters aligned to established subsets identified in healthy and SSc skin (*14, 20*) (**Fig. 2E, S2A-S2C**) and were enriched in expression of lineage-defining genes (**Fig. S2D**).

**Figure 2.**
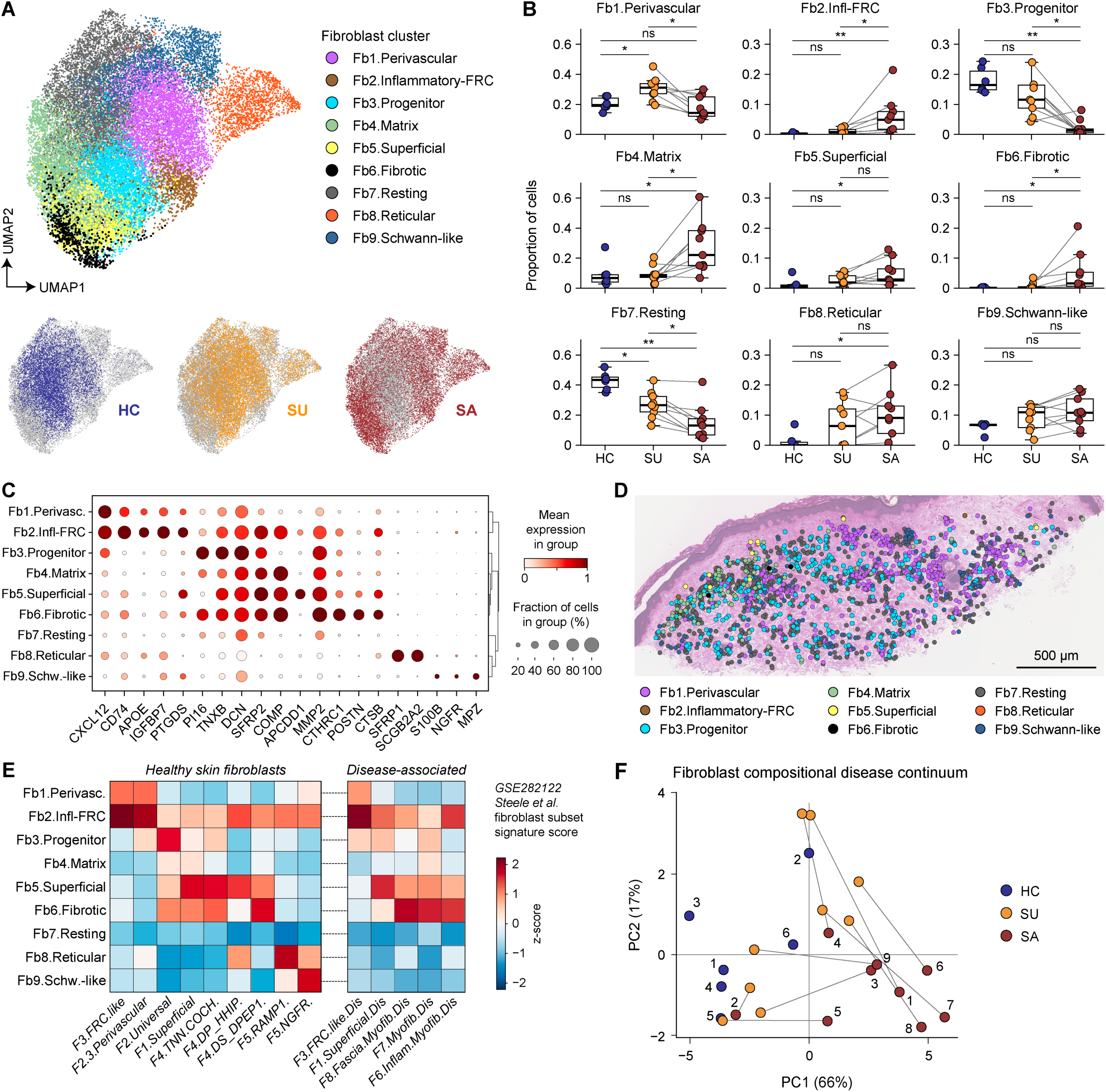
Progenitor fibroblast loss in clinically uninvolved lcSSc skin occurs absent myofibroblast expansion. (**A**) UMAP of 27,000 curated dermal fibroblasts (Fb1-Fb9), split by group. (**B**) Per-donor proportion of each subpopulation, paired SU-SA samples connected. (**C**) Dot plot of subpopulation markers with dendrogram. (**D**) Representative section with fibroblasts overlaid on H&E image, colored as in (A). (**E**) Steele et al. reference fibroblast signature enrichment (z-score; GSE282122) across Fb1-Fb9, split into healthy and disease-associated signatures. (**F**) PCA of per-donor fibroblast composition (PC1, 66%; PC2, 17%), paired SU-SA connected. Box plots, median and IQR; paired SU-SA connected. HC comparisons, two-sided Mann-Whitney U test; SU vs SA, two-sided Wilcoxon signed-rank test on donor pairs; Benjamini-Hochberg correction; *p<.05, **p<.01. Scale bar, 500 µm.

LGR5^+^SCARA5^+^PI16^+^ fibroblasts are a stem-like population enriched in healthy skin and progressively reduced during SSc, but not in other fibrotic conditions such as localized scleroderma or chronic graft-versus-host disease (*14, 16, 20*). Neither their functional relevance to disease nor their status in clinically uninvolved skin has been characterized. Fb3.Progenitor cells were distributed throughout the dermal interstitium, between perivascular adventitial regions (**Fig. 2D**). Transcriptionally, they aligned to the Universal and Fibro-LGR5 populations (**Fig. 2E, S2A-S2C**), preferentially expressing stem/universal-lineage genes (e.g., CD34, CD55, LGR5, PI16, SCARA5, SFRP2, WIF1) (**Fig. S2D**). Fb3.Progenitor cells were enriched in HC and reduced in both SU and paired SA skin, albeit with high donor-to-donor variability (**Fig. 2A, 2B**). We complemented these per-donor comparisons with scCODA, a Bayesian model that jointly evaluates cell-type proportions to account for their compositional interdependence (*28*). This analysis indicated a credible reduction of Fb3 and Fb7 cells already in clinically uninvolved skin relative to healthy controls, with a further decrease in SA (**Fig. S2E**). Expansion of inflammatory, matrix-producing, and fibrotic subsets (Fb2, Fb4, Fb6) was apparent only in SA.

To summarize these coordinated proportional shifts, we performed principal-component analysis (PCA) on fibroblast subset proportions. Samples showed an ordered trend, HC-to-SU-to-SA, establishing clinically uninvolved skin as compositionally intermediate between healthy and fibrotic tissue (**Fig. 2F**). Variance partitioning showed that Fb3.Progenitor abundance accounted for the largest share of compositional variation by study group (**Fig. S2F**). Thus, homeostatic fibroblast loss is a distinguishing subclinical feature of lcSSc.

### Progressive rewiring of homeostatic fibroblast gene programs is evident in clinically uninvolved skin

To define disease-associated fibroblast transcriptional programs we performed differential gene expression analysis on pseudo-bulked fibroblasts across study groups. We filtered for genes that were (i) significantly altered between HC and SA, and (ii) intermediate in SU, yielding a 655-gene set that distinguished the three study groups (**Fig. 3A**; top 159 at FDR < 0.01 shown). Pathway analysis identified terms enriched in HC fibroblasts for skeletal morphogenesis, anterior-posterior pattern specification, and elastic fiber formation, whereas terms enriched in lcSSc included ECM organization, TNF-α/NF-κB signaling, mesenchymal cell proliferation, smooth-muscle contraction, angiogenesis, and Wnt antagonism (**Fig. S3A**).

**Figure 3.**
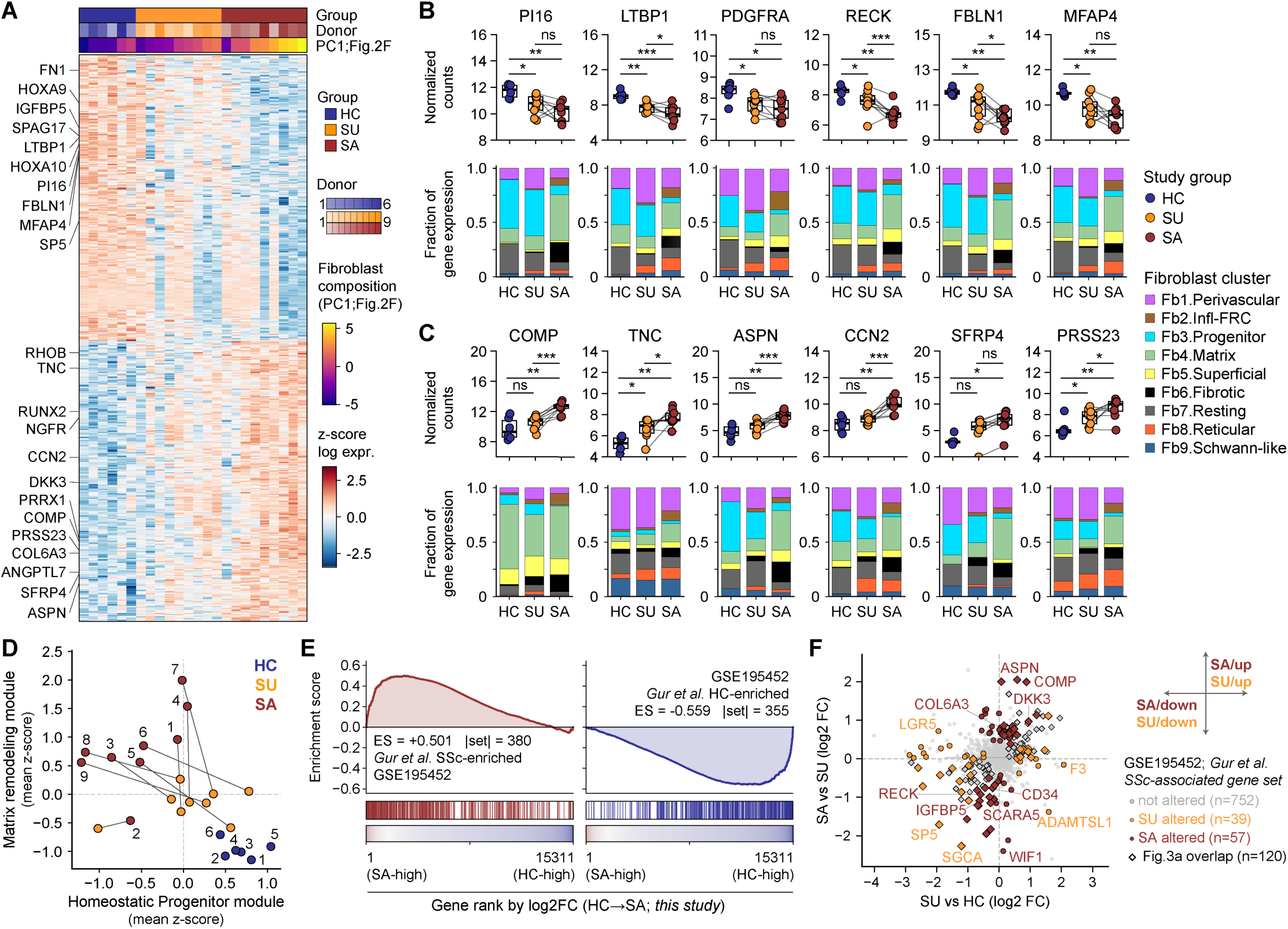
Homeostatic fibroblast transcriptional programs decline in clinically uninvolved skin alongside partial engagement of matrix remodeling genes. (**A**) Heat map of the top 159 genes (FDR < 0.01; 84 down, 75 up) from the 655-gene HC-to-SA discovery set (FDR < 0.05) across samples annotated by group, donor, and PC1 from Fig. 2F; selected genes labeled. (**B** and **C**) Pseudo-bulk expression (log2 TMM-CPM) of representative down-regulated (B: PI16, LTBP1, PDGFRA, RECK, FBLN1, MFAP4) and up-regulated (C: COMP, TNC, ASPN, CCN2, SFRP4, PRSS23) genes by group, with fibroblast-subpopulation contribution below. (**D**) Per-sample homeostatic-progenitor versus matrix-remodeling module scores (genes in Fig. S3C); numbers denote donors, paired SU-SA connected. (**E**) GSEA of Gur et al. (GSE195452) SSc-enriched (left; 380 genes) and HC-enriched (right; 355 genes) gene sets against all 15,311 Visium HD fibroblast genes ranked by HC-to-SA log2 fold change; ES, enrichment score. (**F**) Per-gene SU-vs-HC versus SA-vs-SU log2 fold change for Gur et al. SSc-associated genes, classified as altered in SU, altered in SA, or not altered; diamonds mark genes also in (A). Box plots, median and IQR; paired SU-SA connected; *FDR<.05, **FDR<.01, ***FDR<.001.

Dermal fibroblasts underwent a progressive, graded transcriptional disruption organized along two opposing axes. Reduction of homeostatic genes was already evident in SU, and many were further lost in SA (**Fig. 3A**, **3B**), including markers of universal/progenitor identity or structural cell maintenance (e.g., PI16, PDGFRA, SPAG17, MFAP4, SP5 (*14, 19, 20, 23, 29*)) and those involved in restraining matrix turnover, latent TGF-β activation, and elastic fiber formation (e.g., LTBP1, RECK, FBLN1 (*30, 31*)). The expression of multiple homeobox genes (e.g., HOXA10, HOXA9, HOXD8) were also disrupted in SU (**Fig. S3B**). This family of genes encodes positional information during embryonic development, directs fibroblasts to anatomically distinct skin regions, and sustains adult stromal cell function (*32–34*). We also observed a progressive induction of the myofibroblast developmental and functional program (e.g., COMP, TNC, ASPN, CCN2, SFRP4, PRSS23 (*14, 16–18, 20*)) largely restricted to SA (**Fig. 3A**, **3C**), with TNC and PRSS23 expression already induced in SU.

To capture the transcriptional shift within SSc-associated fibroblasts, we focused on Fb3.Progenitor (HC-enriched) and Fb4.Matrix (SA-enriched) clusters, which span the homeostatic and remodeling ends of the fibrotic continuum. The top 20 up- and 20 down-regulated genes ranked by HC-vs-SA effect size were used to form *homeostatic* and *matrix-remodeling* modules (**Fig. S3C**). Per-sample module scoring showed the transcriptionally intermediate state of SU, with fibrosis-associated cell reprogramming already underway (**Fig. 3D**). This homeostatic-to-remodeling shift is consistent with a cell trajectory analysis showing stem-like SCARA5^+^PI16^+^ fibroblast reprogramming into pathogenic POSTN^+^PRSS23^+^COMP^+^ myofibroblasts during SSc (*16*). Genes defining this progenitor fibroblast trajectory separated our samples by study group, consistent with an inferred myofibroblast differentiation axis (**Fig. S3D**).

For independent validation of this transcriptional rewiring, we ranked all genes by their HC-vs-SA fold change and tested them by gene-set enrichment analysis (GSEA) against modules derived from a large public scleroderma single-cell skin data set (Gur et al., GSE195452 (*14*)) (**Fig. 3E**). GSEA revealed concordance between the two data sets for both HC- and SSc-associated features. Individual gene expression patterns revealed the components of SU homeostatic destabilization (e.g., LGR5, F3, SP5) that occurred independently of the SA fibrosis signature (e.g., *induced*: ASPN, COMP, DKK3; *reduced*: RECK, IGFBP5, WIF1) (**Fig. 3F**).

Collapse of the Fibro_LGR5 progenitor cells in Gur et al. was inversely correlated with our remodeling module (**Fig. S3E**), while our homeostatic and remodeling modules distinguished the Gur et al. study groups: Healthy, lcSSc-affected, and dcSSc-affected skin (**Fig. S3F**). Thus, fibroblast compositional and transcriptional changes occurred in parallel, evident in SU, implicating clinically uninvolved skin as a distinct, subclinical disease state.

### Fibroblast cell- and gene-level positional organization is partially eroded during lcSSc

In healthy dermis, transcriptionally distinct fibroblast populations occupy defined positions that shape their contributions to wound repair, and this organization is disrupted in fibrotic disease (*20, 35–37*). To determine whether SSc-associated compositional and transcriptional changes were accompanied by spatial reorganization, we measured the distance between each fibroblast and the dermal-epidermal junction (DEJ), a defining polarity axis of human skin (**Fig. 4A, 4B, S4A**). Fibroblast subsets segregated by proximity to anatomical features (e.g., vasculature, adnexal, DEJ): Fb5.Superficial papillary fibroblasts nearest the DEJ; Fb8.Reticular fibroblasts in the deep reticular dermis; Fb3.Progenitor and Fb7.Resting fibroblasts in the intervening dermis (**Fig. 4C, S4B**). SSc-altered populations (e.g., Fb3, Fb4) showed a significant DEJ-depth change between SA and HC, with an intermediate trend for SU (**Fig. 4D**).

**Figure 4.**
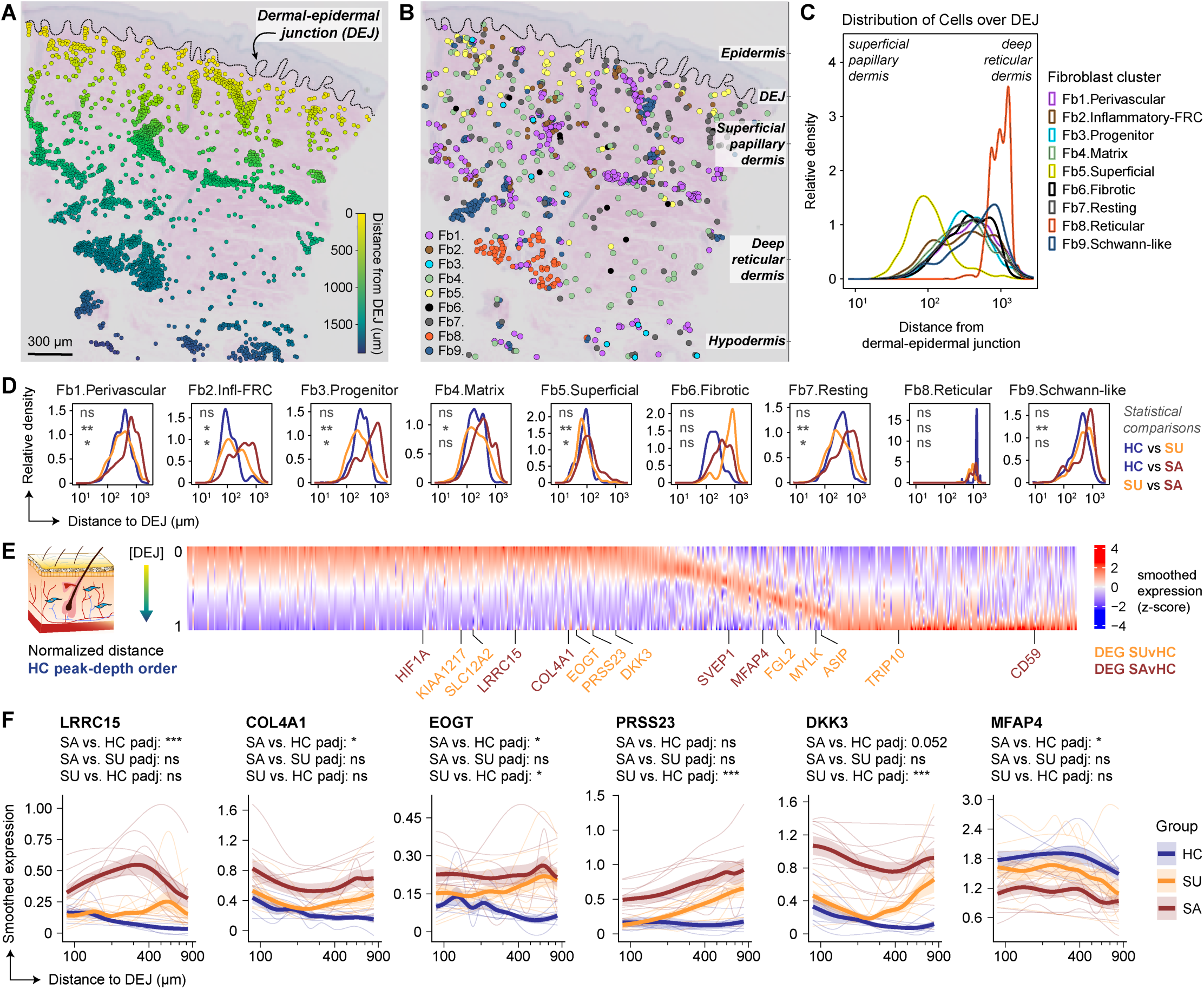
Fibroblast cell- and gene-level positioning along the dermal depth axis is partially eroded in lcSSc with changes evident in clinically uninvolved skin. (**A**) Representative section with all dermal cells colored by distance from the dermal-epidermal junction (DEJ; µm), with the DEJ traced. (**B**) The same section colored by Fb1-Fb9, with dermal zones labeled. (**C**) Relative density of each subpopulation versus DEJ distance, pooled across samples. (**D**) Per-subpopulation DEJ-distance density by group, with the three pairwise comparisons indicated. (**E**) Smoothed expression (z-score) of the 655 disease-axis genes across normalized DEJ distance in HC, genes ordered by peak-expression depth in HC; the 15 genes with a significant between-group difference in depth profile (sample-aware, generalized additive model, FDR ≤ 0.1) are marked (orange, SU vs HC; red, SA vs HC). (**F**) Smoothed expression versus DEJ distance for six representative genes (LRRC15, COL4A1, EOGT, PRSS23, DKK3, MFAP4) by group (thin lines, per sample; bold line, group mean; shaded, SEM); adjusted p values shown. *p<.05, **p<.01, ***p<.001 (D: Mann-Whitney U, Benjamini-Hochberg).

To test whether SSc-fibroblast spatial features extended to the transcript level, we ordered the 655-gene discovery set by peak expression depth in HC dermis and examined if the DEJ distance of each was preserved in SU or SA. We identified 15 genes with statistically significant positional shifts along the DEJ axis relative to HC (**Fig. 4E**). Positionally altered genes were related to matrix biology and myofibroblast function, including those repositioned in SU (EOGT, PRSS23, DKK3) or only in SA (LRRC15, COL4A1, MFAP4) (**Fig. 4F**). Thus, fibroblast positional organization was partially eroded at both the transcript and cluster levels, with some changes present in clinically uninvolved skin and others restricted to affected tissue.

### Dermal vascular smooth muscle cells lose their contractile identity and acquire a synthetic, activated state during lcSSc

Vasculopathy and Raynaud’s phenomenon – reactive vasospasm in the distal digits – are defining early features of scleroderma that precede fibrosis (*13*), yet the spatial and transcriptional perturbations to the dermal vascular compartment during SSc are poorly understood. Independent clustering of smooth muscle and endothelial cells from Visium HD data resolved six transcriptionally distinct populations: three vascular smooth-muscle cell subsets (vSMC-1, vSMC-2, vSMC-3), two vascular endothelial cell subsets (VEC-1, VEC-2), and one lymphatic endothelial cell subset (LEC). Spatial profiling resolved a seventh, non-vascular subset (non-vSMC) that overlapped transcriptionally with vSMC-1 (**Fig. 5A**). The vSMC-1 subset was progressively displaced by vSMC-3 in lcSSc (**Fig. 5B, S5A**). VEC and LEC were characterized but unchanged across study groups (**Fig. S5A-S5C**).

**Figure 5.**
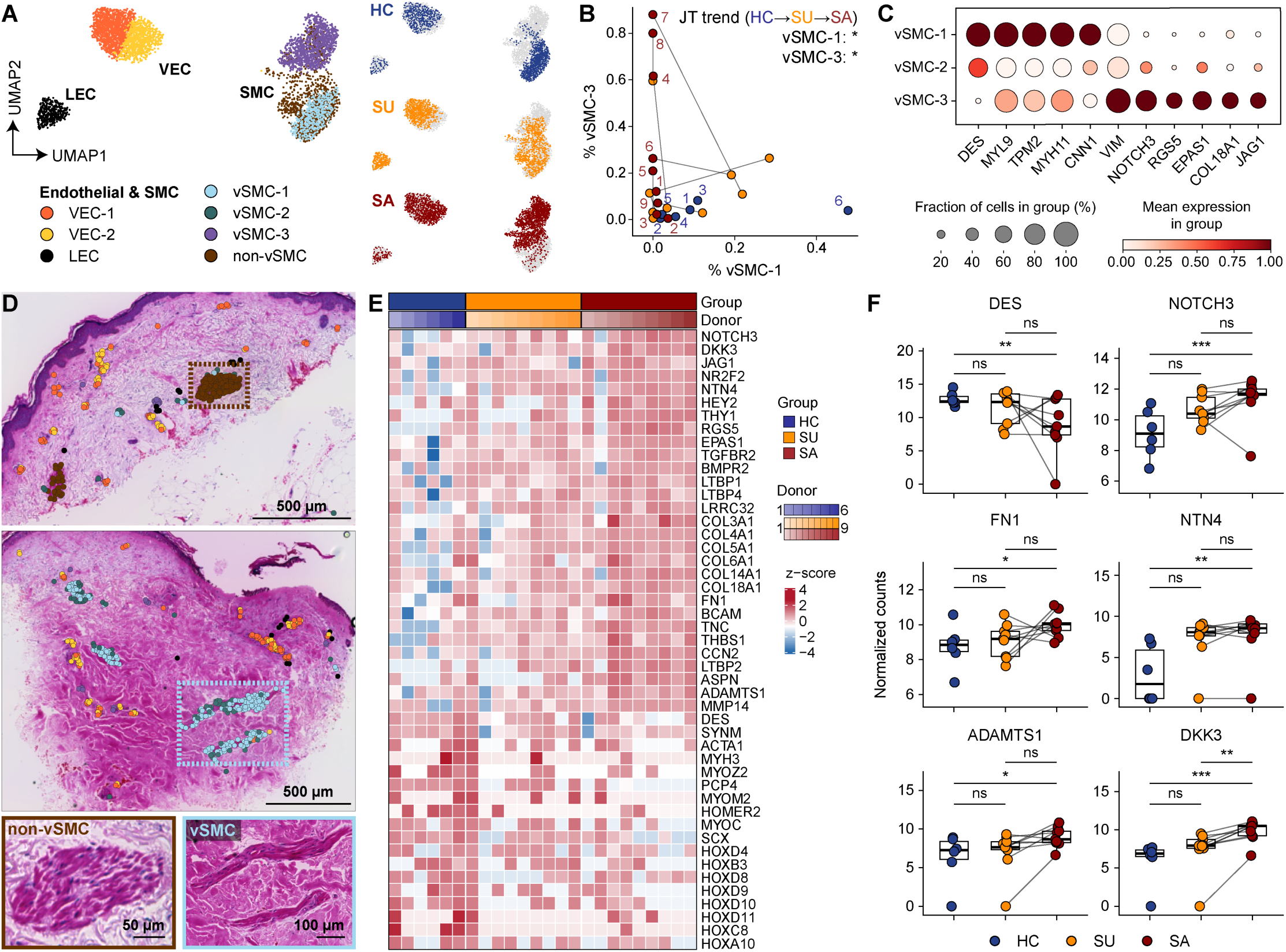
Dermal vascular smooth muscle shifts from a contractile state toward a synthetic, activated state during lcSSc. (**A**) UMAP of the dermal endothelial and smooth-muscle compartment (vSMC-1/2/3, non-vSMC, VEC-1/2, LEC), shown overall and split by group; non-vSMC is a vSMC-1-like population localized to arrector pili. (**B**) Per-sample %vSMC-1 versus %vSMC-3; numbers denote donors, paired SU-SA connected (Jonckheere-Terpstra trend across HC-SU-SA: vSMC-1 p=.012, vSMC-3 p=.016). (**C**) Dot plot of vSMC-1/2/3 markers. (**D**) Representative sections with vascular cells overlaid on H&E image, with H&E insets of a non-vSMC (arrector pili; brown) and a vSMC-enriched vessel (blue). (**E**) Pseudo-bulk heat map (z-score) of vascular identity, remodeling, and positional (HOX) genes by sample and group. (**F**) Per-sample pseudo-bulk expression of DES, NOTCH3, FN1, NTN4, ADAMTS1, and DKK3 by group. Box plots, median and IQR; paired SU-SA connected; *FDR<.05, **FDR<.01, ***FDR<.001 (F: limma-voom contrasts). Scale bars, 500 µm (D), 50/100 µm (insets).

The HC-enriched vSMC-1 subset expressed a contractile program (DES, MYL9, TPM2, MYH11, CNN1) whereas the SA-enriched vSMC-3 expressed a synthetic, activated program (VIM, NOTCH3, EPAS1, COL18A1, JAG1) (**Fig. 5C, S5B**). Non-vSMC were transcriptionally similar to vSMC-1 but spatially restricted to the arrector pili – smooth muscle bundles attached to the hair follicles (**Fig. 5D**) – and showed no disease-associated changes. Pseudo-bulk analysis of the vSMC revealed progressive induction of a vascular remodeling and matricellular program (NOTCH3, DKK3, NTN4, FN1, ADAMTS1) that increased across the HC-to-SA continuum with intermediate trends in SU (**Fig. 5E**, **5F**). Expression of anterior-posterior positional genes (HOX family) was enriched in the vSMC compartment of HC and trended lower in both SU and SA, paralleling our observations in fibroblasts.

Few studies have profiled the vascular compartment during fibrotic disease, in part because these cells are technically challenging to recover. Some of the vSMC changes we observed in lcSSc mirrored those reported in localized scleroderma (GSE264508 (*38*); **Fig. S5D**), including induction of vascular remodeling (JAG1, NOTCH3, NTN4) and loss of contractile or structural muscle genes (DES, SYNM, ACTA1). Together, these findings indicate that dermal vascular smooth muscle undergoes a contractile-to-synthetic state transition that parallels the fibroblast state changes of lcSSc described above.

### Collapse of a homeostatic progenitor fibroblast niche is a defining feature of clinically uninvolved skin

To examine fibroblast-centered dermal organization, we characterized spatial neighborhoods defined by the cell type composition within a 30 μm radius of each fibroblast, which captured cell-cell contacts and short-range interactions (**Fig. 6A**) (*39*). Unsupervised clustering of proximal cell networks identified seven fibroblast-anchored niches (Ni1-Ni7), each defined by distinct cell composition and spatial position: (Ni1) Homeostatic fibroblast-enriched; (Ni2) Macrophage-enriched; (Ni3) T cell-enriched; (Ni4) Diffuse, low-structure, disease-enriched stroma; (Ni5) Smooth muscle-enriched, vascular-associated; (Ni6) Adnexa-enriched; (Ni7) Vascular/endothelial-enriched (**Fig. 6B, S6A**).

**Figure 6.**
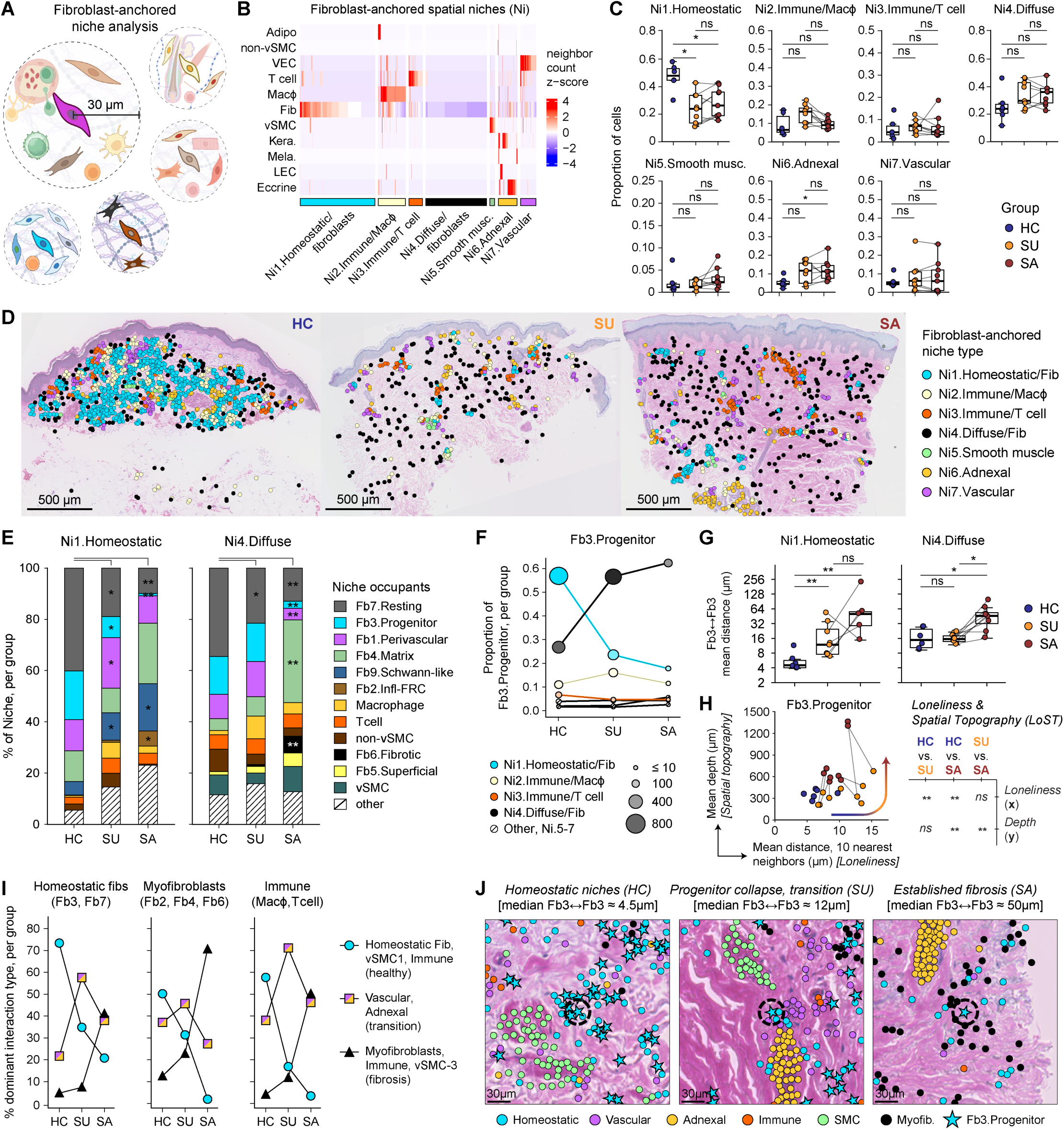
Homeostatic niche collapse and progenitor fibroblast sparsity are evident in clinically uninvolved skin. (**A**) Schematic of fibroblast-anchored niche analysis (30 µm neighborhoods). (**B**) Neighbor-composition heat map (z-score) defining seven niches (Ni1-Ni7); columns, individual fibroblasts; rows, neighbor cell types. (**C**) Per-sample niche occupancy by group. (**D**) Representative sections colored by niche for HC, SU, and SA. (**E**) Composition of the homeostatic (Ni1) and diffuse (Ni4) niches per group (asterisks, significant change versus HC). (**F**) Fb3.Progenitor distribution across niches per group (marker size, cell count). (**G**) Mean distance from each Fb3.Progenitor to its nearest Fb3.Progenitor within Ni1 and Ni4 by group. (**H**) Per-sample Fb3.Progenitor LoST analysis (x, loneliness: mean distance to the 10 nearest cells; y, spatial topography: mean distance from the DEJ), with significance of each component beside the plot. (**I**) Dominant neighboring cell class within 30 µm of homeostatic fibroblasts (Fb3, Fb7; left), myofibroblasts (Fb2, Fb4, Fb6; middle), and immune cells (macrophages, T cells; right), shown as the percentage of cells per group whose dominant neighbors belong to a healthy-associated (homeostatic fibroblast, vSMC-1, immune), transitional (vascular, adnexal), or fibrosis-associated (myofibroblast, immune, vSMC-3) class. (**J**) Representative H&E crops per group with Fb3.Progenitors (stars) and neighboring cell types (median Fb3-to-Fb3 distance shown). Box plots, median and IQR; paired SU-SA connected; HC comparisons, two-sided Mann-Whitney U test; SU vs SA, two-sided Wilcoxon signed-rank test on donor pairs; Benjamini-Hochberg correction; *p<.05, **p<.01. Scale bars, 500 µm (D), 30 µm (J).

Ni1.Homeostatic was enriched in HC and reduced in SU and SA, the collapse of a defining fibroblast interaction hub of healthy dermis (**Fig. 6B**, **6C**). Ni1 contained mostly Fb7.Resting and Fb3.Progenitor cells in HC, which were broadly distributed throughout the dermis (**Fig. 6D, S6B**). The surviving Ni1 nodes in SU had an altered cell composition – Fb7 and Fb3 were replaced by Fb1 and Fb9. Reciprocally, Ni4 abundance trended upward in lcSSc donor tissue and was the principal pathogenic niche containing Fb4.Matrix and Fb6.Fibrotic cells in SA (**Fig. 6E**). Notably, the Fb3.Progenitor reduction in SU was niche-specific, occurring *only* within Ni1 and leaving Ni4 as the dominant Fb3 reservoir (**Fig. 6F**). This likely indicated repositioning between niches, as reflected by changes in the local tissue density of LGR5^+^SCARA5^+^PI16^+^ fibroblasts during lcSSc. In HC, Fb3 of Ni1 were in very close contact with one another, on average ∼4.5 μm apart, which increased to ∼12 μm in SU, and ∼50 μm in SA (**Fig. 6G**). By comparison, Fb3 in Ni4 were more dispersed, even in HC. To examine how altered homotypic progenitor fibroblast networks related to distance from the DEJ, we calculated a ‘<u>Lo</u>neliness-and-<u>S</u>patial-<u>T</u>opography’ (LoST) score. This custom metric emphasized group-level changes in progenitor fibroblast spatial biology: Fb3 were isolated from their homotypic network in SU but only repositioned to the deeper reticular dermis in paired-donor SA samples (**Fig. 6H**).

Dysfunctional communication among vascular-immune-stromal cell networks is a feature of fibrotic diseases such as SSc (*19, 24, 40–42*). Besides the homotypic interactions of Fb3 and Fb7, their dominant neighbors in HC skin were vSMC-1, macrophages, or T cells. By comparison they were most proximal to vascular or adnexal cells in SU, and myofibroblast, immune, or vSMC-3 in SA (**Fig. 6I, 6J, S6C**). Fb3 gene expression changes also tracked with study group (HC vs. SU), more than niche occupancy (Ni1 vs Ni4) (**Fig. S6D**). To examine the altered Fb3 communication networks of lcSSc, we used CellChat to infer signaling relationships that change between HC and SA. In lcSSc, we found an increase in the putative level of fibroblast signaling through matrix components (e.g., COLLAGEN) and of macrophage migration inhibitory factor (MIF) pathway signaling (**Fig. S6E**). Thrombospondin (THBS) signaling, associated with pathogenic matrix remodeling and latent TGF-β activation (*43, 44*), had the strongest inferred sender signal coming from Fb2.Inflammatory-FRC, Fb4.Matrix, and Fb6.Fibrotic cells in SA (**Fig. S6F**).

Thus, the cellular and transcriptional changes of the lcSSc dermis occur alongside collapse of the homeostatic stromal niche that houses LGR5^+^SCARA5^+^PI16^+^ fibroblasts in healthy skin. Collapse of this niche and rewiring of homeostatic stromal interaction networks are already evident in clinically uninvolved lcSSc skin.

### A composite spatial-molecular index orders samples along a clinical continuum distinguished by homeostatic and fibrotic features

To integrate the compositional, transcriptional, and positional changes observed across our cohort into a single unifying framework, we assembled a per-sample feature matrix spanning major disease-associated phenotypes. We computed pairwise Spearman correlations across all study groups and clustered the resulting feature-by-feature matrix (**Fig. 7A**). Clustering resolved two coherent, anti-correlated blocks: (i) a fibrotic block of disease-expanded populations (Fb2, Fb6, vSMC-3), SSc-associated transcriptional signatures, Fb3 positional changes, and clinical severity; and (ii) a healthy tissue block of HC-enriched populations (Fb3, vSMC-1), homeostatic/stem-like fibroblast modules, and Ni1 abundance. Donor age and disease duration showed no strong association with either block. PCA incorporating study-wide compositional, transcriptional, and spatial features ordered samples largely by group (HC-to-SU-to-SA) along a dominant axis, PC1 (**Fig. 7B**).

**Figure 7.**
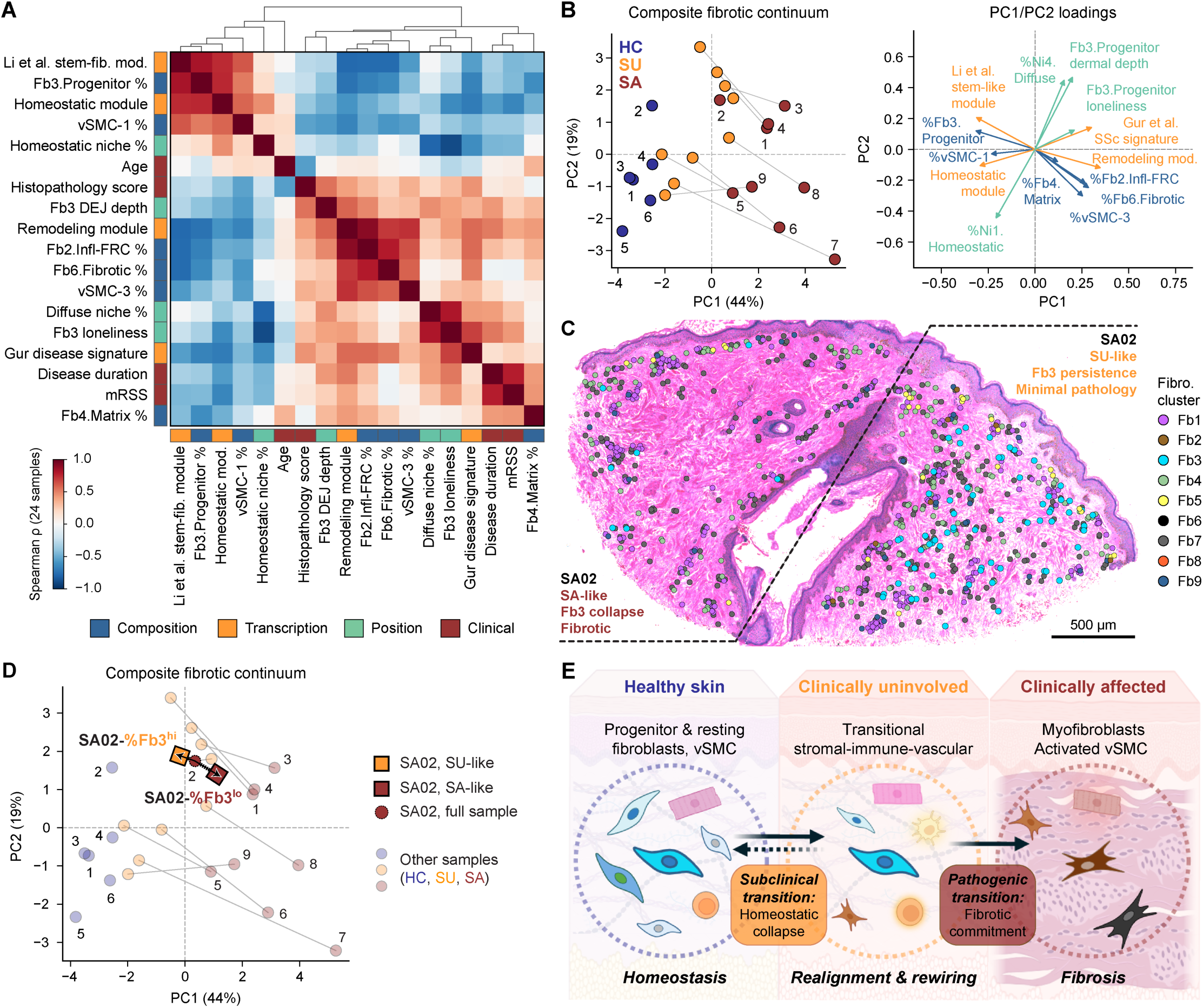
A composite spatial-molecular index orders samples along a continuum defined by anti-correlated blocks of homeostatic and fibrotic features. (**A**) Spearman correlation heat map of 18 per-sample features (composition, transcription, position, clinical), hierarchically clustered; bars indicate feature domain. (**B**) PCA of the 24 samples on the 14 non-clinical features (left; PC1, 44%; PC2, 19%; paired SU-SA connected) with PC1/PC2 loadings (right). (**C**) SA02 tissue section with fibroblasts (Fb1-Fb9) overlaid on H&E image; the dashed line separates a progenitor-rich, near-normal (SU-like) region from a progenitor-depleted, fibrotic (SA-like) region. Scale bar, 500 µm. (**D**) Projection of the full SA02 section and its two in silico halves (SA02-%Fb3^hi^ and SA02-%Fb3^lo^) onto the PCA in (B); other samples faded. (**E**) Model of clinical transition states along the fibrotic continuum of scleroderma. *Subclinical transition* (HC-to-SU): homeostatic neighborhood collapse, loss of progenitor fibroblasts and a trend toward fewer contractile vSMC. *Pathogenic transition* (SU-to-SA): myofibroblast and activated vSMC expansion, fibrosis. Fb1.Perivascular; Fb2.Inflammatory-FRC; Fb3.Progenitor; Fb4.Matrix; Fb5.Superficial; Fb6.Fibrotic; Fb7.Resting; Fb8.Reticular; Fb9.Schwann-like.

In one clinically affected biopsy (SA02, forearm), we observed a striking within-sample histological demarcation that permitted an illustration of our clinical disease continuum. One half of SA02 had minimal pathology and retained normal fibroblast cellularity, including an intact Fb3.Progenitor compartment, while the other half was fibrotic and Fb3-depleted (**Fig. 7C**). We digitally bisected SA02 into two pieces: “SU-like” (histologically normal, %Fb3^hi^) and “SA-like” (fibrotic, %Fb3^lo^). The SA02-%Fb3^hi^ half aligned with SU group averages for fibroblast cellular and transcriptional metrics (**Fig. S7A**). While the %Fb3^lo^ half matched other SA samples on progenitor collapse and homeostatic module scores, it did not show the high level of myofibroblast expansion, remodeling signature, or vascular changes of other SA samples, consistent with progenitor loss as a subclinical perturbation in lcSSc. Remarkably, these two *in silico*-derived halves projected in opposite directions from the full SA02 sample, towards other SU or SA samples, respectively (**Fig. 7D**).

These data define distinct spatial-molecular states of lcSSc skin relative to healthy tissue, in which LGR5^+^SCARA5^+^PI16^+^ progenitor fibroblasts occupy dense stromal niches. Clinically uninvolved SU skin is defined by a *subclinical state* of tissue rewiring: (i) reduction of LGR5^+^SCARA5^+^PI16^+^ fibroblasts; (ii) a trend toward fewer vSMC-1; (iii) loss of stem-like, positional, and contractile gene programs; and (iv) homeostatic fibroblast niche collapse, driving changes to the stromal-immune-vascular network of healthy skin. Clinically affected SA skin is defined by an *overt fibrotic state*: (i) pathogenic myofibroblast and synthetic vSMC-3 expansion; (ii) induction of the canonical matrix biology gene programs of fibrosis; and (iii) deep dermal isolation of rewired fibroblast populations (**Fig. 7E**). Collectively, these findings identify clinically uninvolved lcSSc skin as a biologically distinct state intermediate between healthy and fibrotic skin, and an integrated spatial-molecular index orders samples along this continuum by their homeostatic and fibrotic features.

## DISCUSSION

In this study we characterized the transcriptional and spatial remodeling of dermal fibroblasts across healthy skin from control donors and clinically uninvolved or affected skin in patients with lcSSc. Our central finding is that clinically uninvolved skin, while histologically indistinguishable from site-matched healthy tissue, is biologically distinct yet not overtly fibrotic. It retains some features of healthy tissue while exhibiting changes to the dermal vasculature and stroma, representing a subclinical spatial-molecular disease state that previous analyses could not resolve (*25–27*).

We propose a two-state model that aligns our study with the current understanding of SSc pathogenesis. Clinically uninvolved skin manifests a *subclinical disease state* defined primarily by collapse of a homeostatic fibroblast niche and loss of SSc-associated LGR5^+^SCARA5^+^PI16^+^ cells. These features may facilitate diagnosis in equivocal biopsies from patients with early or suspected disease and could guide preventive strategies to restore homeostasis and prevent irreversible fibrosis. Fibrotic skin marks an *overt disease state* defined by homeostatic cell collapse, the expansion of disease-associated myofibroblasts, and the development of activated vascular smooth muscle. These features were integrated into a spatial-molecular index of disease that segregated samples by study group, HC-to-SU-to-SA.

Progression of fibrosis in established lcSSc is slow; skin scores change little over time (*45*). In patients with very early SSc, more than half meet classification within five years (*13*), while repeat sampling of skin shows a largely consistent molecular phenotype over time (*46*). Yet established lcSSc is not clinically quiescent: 22% of patients develop clinically significant pulmonary fibrosis over 15 years of follow-up (*47*), and calcinosis remains frequent in established disease (*48*). While the median disease duration in our cohort was 14 years from diagnosis, the subclinical state we observed was already present in two early lcSSc donors (<5 years from diagnosis), suggesting it is not merely a feature of established disease. Previous microarray profiling of clinically uninvolved forearm skin biopsies from patients with established lcSSc showed that such tissue can be molecularly abnormal, with two-thirds of patients up-regulating extracellular matrix, TGF-β response, and cardiovascular development genes (*26*). Our study now identifies the cells and structures that underlie this subclinical state: depletion and repositioning of LGR5^+^SCARA5^+^PI16^+^ progenitor fibroblasts, collapse of the homeostatic fibroblast niche, and a trend toward vascular smooth muscle reprogramming, all in clinically and histologically normal skin. Organizational changes occurred in both uninvolved and affected skin, while myofibroblast expansion and fibrotic matrix programs were confined to affected sites. This raises a central question: what are the missing triggers in clinically uninvolved skin that drive vascular or fibrotic pathology at affected sites? One possibility is that patients with established lcSSc no longer harbor the active signals that precipitate additional pathology. Profiling these subclinical features in cohorts of less established disease, including very early SSc and recent onset lcSSc and dcSSc, will be required to define how this state relates to disease progression.

Whether niche reorganization and positional realignment of the dermis in SSc is dependent on intrinsic or extrinsic factors remains an open question. Lineage tracing in mouse skin shows that layer-specific fibroblast programs are largely cell-intrinsic and dictate each population’s contribution to repair (*35*). In humans, anatomically specified, epigenetically maintained expression of homeobox/HOX transcription factors, which encode positional identity in fetal development, are imprinted postnatally in anatomically distinct dermal fibroblasts (*34, 49*) and govern adult stem-cell function (*32*). Differential HOX usage in healthy adult skin tunes fibroblast mechanotransduction during the response to injury, and in hypertrophic scar and keloid formation (*50*). SU fibroblasts and vSMC showed altered HOX gene expression, relative to site-matched HC populations, suggesting that broad changes to stromal anterior-posterior positional programs are a defining subclinical feature of lcSSc, but are not sufficient for fibrosis. One intriguing possibility is that changes to the anatomically imprinted dermal HOX gene program could explain differences in the rate or sites of scleroderma progression, or the distinct origins and outcomes of lcSSc versus dcSSc. Future analysis of lcSSc susceptible versus spared anatomical regions (e.g., forearm versus trunk) would offer a unique opportunity to test this hypothesis.

In this study, we showed that reprogramming of the dermal compartment in lcSSc included a shift of vascular smooth muscle from the contractile phenotype of healthy skin toward a synthetic, activated state, defined by induction of NOTCH and matrix-associated programs. Endothelial NOTCH signaling regulates vascular remodeling during cutaneous repair (*51*), and in SSc, dermal vascular cells contribute to extracellular matrix deposition through endothelial-to-mesenchymal transition (*11*). Consistent with a vascular contribution to disease, multi-modal profiling of skin from recently diagnosed treatment-naïve SSc subjects implicated macrophage-fibroblast interactions at inflamed vascular niches as a key pathogenic process (*52*). Because vasculopathy is an early clinical manifestation of SSc that contributes to fibrosis (*53*), this rewiring of vascular smooth muscle and progenitor fibroblasts suggests that loss of dermal cell homeostasis in SU is a multi-compartment process that could render the tissue permissive to fibrotic commitment.

A naturally occurring histologic border within an affected forearm biopsy (SA02) illustrated the observation of an advancing pathogenic “front” at the periphery of overtly fibrotic areas, one that tends to spread proximally toward unaffected skin during lcSSc progression. Remarkably, the minimal-pathology/SU-like and adjacent fibrotic/SA-like halves of SA02 could be distinguished using only Fb3.Progenitor abundance. Depletion of the LGR5^+^SCARA5^+^PI16^+^ population alongside evident fibrosis suggests that progenitor cell abundance may distinguish the subclinical and pathogenic states. Current criteria for the very early diagnosis of SSc stratify at-risk patients with Raynaud’s phenomenon using serology, nailfold capillaroscopy, and puffy fingers, but include no tissue measures (*13*). Whether progenitor fibroblast niche collapse is already detectable in such patients, before classification criteria are met, is an important question for future studies to address.

The signals that regulate progenitor fibroblast persistence or differentiation remain undefined, and many candidate mechanisms exist. Interaction with extracellular matrix components is essential for fibroblast function (*54*), and the spatial isolation of LGR5^+^SCARA5^+^PI16^+^ cells in SU is consistent with a role for homotypic cell-cell contacts or paracrine signals in sustaining the progenitor state. Immune-derived cues are another set of candidate regulators with T cells having long been implicated in scleroderma (*55*). We previously showed that cutaneous tissue-resident memory CD4^+^CLA^+^CD103^+^ T cells induce an LGR5-enriched, stem-like program in healthy dermal fibroblasts and that this T cell population is reduced in lcSSc and dcSSc (*56, 57*). TGF-β pathway signaling, a central regulator of wound healing and tissue fibrosis (*58*), supports the function of CD103^+^ resident-memory T cells and regulatory T cells that are dysregulated during SSc (*59*). A recently described CSF1/CSF1R axis governing dermal fibroblast-macrophage equilibrium could likewise regulate progenitor persistence or differentiation (*40*). Discriminating among these possibilities will require precise future studies to dissect both the cellular and molecular dysregulation of the vascular-immune-stromal network.

Our study has limitations. Visium HD provided a genome-wide survey at single-cell resolution, but the transcriptional signal from the most abundant gene targets limited the detection of rare populations (e.g., T cell or macrophage subsets), and available segmentation-based modeling did not allow subcellular transcript localization, limiting receptor-ligand inference. Our cross-sectional, exploratory cohort is underpowered to resolve the effects of sex, race, autoantibody type, treatment, or disease duration. SA biopsies were mostly acral with one forearm sampled, whereas HC and SU biopsies were all from the forearm, so the HC-vs-SU comparison is site-matched, but comparisons to SA are somewhat confounded by anatomic site. However, sampling uninvolved forearm and nearby affected skin from the same donors reflects the typical distal-to-proximal pattern of lcSSc involvement. We did not establish whether homeostatic niche collapse is causal in fibrotic progression or whether any direct lineage relationship exists between Fb3.Progenitor or vSMC-1 and SA-expanded myofibroblast or vSMC-3 populations. Subcellular spatial transcriptomics, multiplexed protein imaging, longitudinal sampling, and functional perturbation studies will be required to define the mechanisms that maintain or destabilize the dermal compartment to define disease tempo and spread.

This study established that clinically uninvolved skin from lcSSc patients, despite lacking fibrotic histology, contained measurable molecular and spatial features of subclinical disease. Our data support a model in which the coordinated loss of homeostatic stromal organization is a subclinical feature, separable from myofibroblast program commitment and fibrosis. Current therapies in SSc, including immunomodulatory and anti-fibrotic agents, act on established pathogenic cell populations (*60*). By showing that molecular and spatial changes are already present in clinically uninvolved skin, our data support the notion that disease state can be used to stratify patients, as proposed elsewhere (*61, 62*). Intervention should aim to preserve or restore progenitor fibroblasts and contractile vascular smooth muscle to prevent commitment to an irreversible fibrotic phenotype.

## MATERIALS AND METHODS

### Study design and participants

The objective of this study was to define the cellular, transcriptional, and spatial changes that distinguish healthy skin, clinically uninvolved skin from patients with lcSSc, and clinically affected skin from the same lcSSc patients. This was a cross-sectional, exploratory study of 24 skin biopsies from 15 individuals: 6 healthy controls (HC; one biopsy each) and 9 lcSSc donors, each contributing paired biopsies of clinically uninvolved skin (SU) and of clinically affected skin (SA) (**Table S1**). Participants were enrolled in the Immune-Mediated Diseases Registry and Repository (IMD-RR): Rheumatic Diseases Translational Research Study, Benaroya Research Institute (Seattle, WA). All lcSSc donors had modified Rodnan skin scores of 2-8 (median 4).

Disease duration at the time of biopsy, measured from diagnosis, ranged from 1 to 39 years (median 14), with two donors (lcSSc 06 and 07) within 5 years of diagnosis. HC and lcSSc groups were age- and sex-matched. Median age was 70 years in HC (range 57-73) and 71 in lcSSc (range 42-77). Details including race, ethnicity, disease duration, and medications at the time of biopsy are listed in Table S1. Participants were assigned to groups by clinical status. No formal power calculation was performed: the cohort size for this rare disease reflected the availability of paired lcSSc samples in the IMD-RR. Dermal fibrosis was graded on H&E sections by a board-certified dermatologist blinded to study group. No samples were excluded.

### Tissue collection

Skin biopsies (3-4 mm) were collected at a single research visit by trained clinical staff under local anesthesia. A board-certified dermatologist selected skin sites at the time of collection. HC and SU biopsies were taken from the forearm. SA biopsies were taken from the finger or forearm (**Table S1**). Biopsies were fixed using a single lot of 10% neutral-buffered formalin for 24 hours, transferred into ethanol and paraffin embedded.

### Ethics

Human study protocols were approved by the Institutional Review Board of Benaroya Research Institute (IRB10059; Seattle, WA). All procedures and sample collection were performed at Virginia Mason Franciscan Health (Seattle, WA) after written informed consent of participants. Samples were provided de-identified in accordance with the HIPAA Privacy Rule. The investigators had no access to personal identifiers.

### Statistical analysis

Statistical tests are noted in each figure legend. Complete statistical details are provided in the Supplementary Materials. Group comparisons used the sample or donor, where stated. HC-vs-SU and HC-vs-SA comparisons of per-sample values used two-sided Mann-Whitney U tests; SU-vs-SA comparisons of paired data used two-sided Wilcoxon signed-rank tests on donor pairs. Multiple testing was controlled by the Benjamini-Hochberg procedure within each contrast and family of tests, as specified in the Supplementary Materials; P values or false-discovery rates (FDR) < 0.05 were considered significant unless stated otherwise. For box plots, the center line is the median, the box spans the interquartile range, and whiskers extend to 1.5 times that range.

## List of Supplementary Materials

Additional Materials and Methods Figures S1-S7

Table S1 References (63–73)

## Acknowledgements

We are grateful to: Kassidy Benoscek-Narag, Sylvia Posso, Thien-Son Nguyen, Marlie Wells, and the Clinical Research Center Registered Nurses in the Benaroya Research Institute (BRI) Center for Interventional Immunology for human subject screening, recruitment, and sample collection; the BRI Cell and Tissue Analysis (RRID:SCR_026327), and Genomics (RRID:SCR_026658) facilities for technical assistance and expertise; the BRI Scientific Writing Group for critical reading of the text. BRI imaging and genomics equipment used in this study were generously supported by the M.J. Murdock Charitable Trust.

## Author contributions

Conceptualization: SL, DEF, JSC, DJC, PAM; Data Curation: AKH, LWT, TP; Formal Analysis: AKH, LWT, TP, PAM; Funding Acquisition: HAD, DJC, PAM; Investigation: AKH, LWT, TP, CLS, PAM; Methodology: CLS, HAD, DJC, PAM; Project administration: PAM; Software: AKH, LWT, TP; Resources: SL, JSC; Supervision: HAD, DJC, PAM; Validation: AKH, LWT, TP, PAM; Visualization: HAD, DJC, PAM; Writing – original draft: PAM; Writing – review and editing: AKH, LWT, TP, CLS, SL, DEF, HAD, DJC, PAM.

## Data, code, and materials availability

Gene expression data generated in this study have been deposited in the Gene Expression Omnibus (GEO) under an accession number that will become available upon final publication. Raw sequencing data are available through the database of Genotypes and Phenotypes (dbGaP) under accession number phs005032.v1. Access is controlled and granted to qualified researchers through the dbGaP authorized access process. The code used for analysis will be made available upon final publication on GitHub at https://github.com/BenaroyaResearch (commit id, to be determined).

