## Supplementary Materials for "Homeostatic fibroblast niche collapse in clinically uninvolved skin defines a subclinical state of systemic sclerosis"

### Additional Materials and Methods:

#### Skin histology, imaging, and dermal fibrosis scoring

Formalin-fixed, paraffin-embedded (FFPE) skin biopsies were sectioned and stained with hematoxylin and eosin (H&E) according to manufacturer instructions for the Visium HD workflow (see below) with the following modifications: Hematoxylin incubation time was increased to 2 minutes and additional washes were included after Eosin incubation (Slides were incubated at room temperature for 1 minute each in 100% then 95% EtOH, followed by two quick washes in deionized water). High resolution H&E sections were imaged before Visium HD CytAssist transfer using an ImageXpress imaging system (Molecular Devices). Images were used for histopathology and as the spatial reference for cell segmentation. Analyses used the original images. Dermal fibrosis was graded from 0 to 5 using an established method *(63)* by a dermatologist blinded to study group. H&E images in figures were white-balance corrected for display only (fixed per-channel gains from a pooled background white point of all images; per-image background correction for Fig. S4B). For the correlation analysis (**Fig. 7A**), mRSS and disease duration were set to HC = 0 for reference group samples.

#### RNA quality assessment

RNA was extracted from FFPE tissue sections using the Qiagen RNeasy FFPE Kit (catalog no. 73504) according to manufacturer protocol and analyzed on an Agilent 4200 TapeStation. RNA integrity was evaluated using DV200, the percentage of RNA fragments longer than 200 nucleotides. All samples exceeded the DV200 threshold of 30% (range for all samples: 57-89%) recommended by 10x Genomics for Visium HD Spatial Gene Expression (**Table S1**).

#### Generation of spatial transcriptomics data using Visium HD

Skin FFPE tissue was sectioned onto NEXTERION® 3-D Hydrogel (H) slides (Schott) at 5-micron thickness after testing neighboring sections for RNA quality by RNA extraction and calculation of a DV200 score, as described. Tissue sections were H&E-stained and imaged as described to generate a high-resolution reference for cell segmentation. After destaining and crosslinking, the sections were processed according to manufacturer instructions, including hybridization to human probe set v2 (18,132 probed genes; 10x Genomics) and transferred to a Visium HD slide, using a Visium CytAssist (10x Genomics). Gene expression libraries were constructed and pooled according to their respective area coverage, as determined by Loupe Browser v8 (10x Genomics). Sequencing was carried out on a NextSeq 2000 sequencer, using NextSeq 2000 XLEAP-SBS flowcells (Illumina).

#### Spatial transcriptomics analysis

Base call files were demultiplexed into FASTQ files (*spaceranger mkfastq*), followed by read alignment (*spaceranger count*) to the human genome (GRCh38, Ensembl 110 annotations) using the Space Ranger (v4.0.1) software (10x Genomics). In addition to the default image processing steps, UMI-based registration algorithm was applied to refine the alignment between UMI density and tissue registration. Cell segmentation was performed with the Space Ranger custom implementation of StarDist. Resulting gene count-by-cell matrices were used for downstream processing with Python (v3.13.9). Quality control of cells was assessed with the following metrics and thresholds: number of genes expressed (> 50/cell), number of transcripts (100-6000 UMIs/cell), proportion of mitochondrial transcripts (<20%/cell), and aggregated bin count (5-100 bins/cell). Genes expressed in less than 3 cells per sample were also excluded. The final data set after all filtering contained 96,865 segmented cells.

UMI counts were normalized and log-transformed, followed by the identification of top 2000 variable genes by variance-stabilizing transformation for dimensionality reduction and Leiden clustering using Scanpy (v1.11.0) *(64)*. Cells were annotated with a comprehensive set of curated identity markers from published studies *(14, 16, 20)*. Serial sub-clustering was leveraged to exclude cells that exhibit ambiguous expression profiles from downstream analysis. Non-vascular and vascular smooth muscle cells were distinguished from each other by masking non-vascular structures such as arrector pili.

Comparison of cell proportions across disease groups was performed by two-sided Mann-Whitney U and Wilcoxon signed-rank tests for comparisons to healthy controls and comparisons between clinically uninvolved and affected samples (to account for donor pairing), respectively. Supplemental analysis with scCODA (dependency of pertpy v1.4.0) was applied to confirm proportional differences in the presence of compositional bias *(28)*. Principal-component analyses of per-sample features were performed with scikit-learn *(65)*.

#### Dermal-epidermal junction distance analysis

Dermal-epidermal junction (DEJ) was manually drawn on high-resolution H&E scans. The position of each cell relative to the DEJ was defined as the minimum distance, in microns, between the cell centroid and the drawn DEJ boundary. To identify genes that characterize the programs associated with the depth of skin, the expression of genes over the distance to DEJ was smoothed within each disease group using Gaussian generalized additive models (GAMs). The modeled range of distance to DEJ was limited to the most restrictive 5%-95% distance range across the three disease groups to prevent overfitting in regions with too few cells. Tests for differential relationship of binarized gene expression (zero or non-zero expression) to distance to DEJ were performed using binomial GAMs with factors accounting for: library depth per cell, overall distance-agnostic difference in expression between disease groups, and variation per sample, using the function from mgcv (v1.9.4):

*bam(expression ~ disease group + s(log distance to DEJ, bs = "cr") + s(log distance to DEJ, bs = "cr", by = disease group) + s(log distance to DEJ, sample, bs = "fs", k = 5) + offset(log library size), family = binomial(link = "cloglog"), method  = "fREML", discrete = TRUE)*

Between group differences were tested as the significance of group-specific difference factor *s(log distance to DEJ, by = disease group)* only on genes expressed in at least 2 donors per group. P-values were adjusted using Benjamini-Hochberg adjustment across the limma-voom 655 discovery genes set defining the HC to SA disease continuum, across all pairwise comparisons.  Other distance features were also computed per cell: the number of adjacent cells within a 30 μm radius of each cell, the log2 distance between each cell and its nearest epidermal keratinocyte, and the log2 distance between each cell and its nearest vascular cell.

#### Spatial niche assignment

For each annotated population, spatial niches were identified using a clustering approach on cell composition within a 30 μm radius of each cell. Louvain clustering with igraph (v2.2.2) *(66)* was performed on a random sample of 6000 cells at varying resolutions from 1.00 to 1.012 in increments of 0.0008. Silhouette coefficient was used to evaluate resulting clusters where the 8-niche solution was chosen for interpretability. Niches 1 and 2 were combined due to similarity in composition and spatial co-occurrence. Cells not included in the clustering were then assigned a spatial niche by identifying their 10 most similar cells among the 6000 clustered cells, and taking the niche most commonly represented among those neighbors. For each fibroblast niche, per-sample proportions in healthy control (HC) were compared to SU and to SA samples using two-sided Mann-Whitney U tests, with Benjamini-Hochberg FDR correction applied across all niches.

#### Pseudobulk comparisons

Pseudobulk transformation was applied to annotated cell populations per donor, followed by linear modeling and differential gene expression analysis across disease groups using Limma in R (v4.4.2) *(67)*. Genes with less than 1 count-per-million (CPM) in at least 10% of samples were excluded. Repeated measures from the same donor were accounted for by modeling donor as a random effect. Benjamini-Hochberg correction was applied to p-values to correct for multiple testing. Heatmaps represent Z-scores of log2-transformed CPMs normalized to the effective library size from TMM normalization computed with edgeR *(68–70)*. Genes were included in the “discovery set” for fibroblasts if they differed significantly between the healthy (HC) and disease (SA) endpoints and showed intermediate behavior in the SU group consistent with a continuum.

#### Inference of cell communication

Cell signaling networks in the annotated spatial data were assessed using CellChat (v2.2.0) *(71)* following the published vignette (https://github.com/sqjin/CellChat). For each sample, the average interaction probability was computed per pathway for any interaction involving fibroblasts as either a sender or receiver. To compare the degree of cell signaling across disease groups, t-tests were applied to log-transformed average interaction probabilities for pathways with a non-zero interaction probability in at least 12 samples.

#### Use of public datasets and gene modules

Gene module scores were computed per cell whereby the mean log-normalized expression of the module genes was compared against that of expression-matched control genes, and the difference was taken as the score. Module genes not detected in the dataset were excluded. Fibroblast subset modules from Steele et al. (GSE282122) were directly downloaded from the published supplemental files *(20)*. Fibroblast modules from Gur et al. (GSE195452) were derived from differential gene expression in their annotated dataset *(14)*. Li et al.

stem-like fibroblast and myofibroblast gene modules were taken directly from figures *(16)*. To examine the disease profile of vascular smooth muscle cells in scleroderma, we performed pseudobulk analysis on extracted, annotated data from Hutchins et al. (GSE264508) *(38)*. The homeostatic progenitor and matrix remodeling modules were derived from the fibroblast discovery gene set whereby the top 20 upregulated and 20 downregulated genes in progenitor and matrix fibroblast populations (Fb3, Fb4) were used.

**Pathway enrichment analysis**

Gene set enrichment analysis (GSEA) of the Gur et al. disease gene sets and the associated barcode plot were generated by ranking genes by HC vs. SA log2 fold change and computing the weighted running enrichment scores and associated P-values of the genes in the sets *(72)*. For over-representation analysis, upregulated and downregulated genes in the fibroblast discovery were tested for over-representation in the MSigDB Hallmark, Reactome and GO Biological Process libraries using Enrichr *(73)* (Fisher exact test against the Enrichr default background). Representative terms are shown as signed -log10 FDR, with selected overlapping discovery genes listed beneath each bar.

**Fibroblast loneliness and spatial topography**

For each Fb3.Progenitor fibroblast, spatial "loneliness" was defined as the mean Euclidean distance to the 10 nearest cells of any type within the same tissue section (nearest neighbors from a KD-tree on spot-grid coordinates converted to μm). For the same cell, "spatial topography" was then defined as the centroid distance to the dermal-epidermal junction (DEJ depth; distances from the DEJ-tracing pipeline). Both metrics were averaged across all Fb3.Progenitor cells per sample and plotted per sample, colored by group with paired SU/SA donors connected. The two components were each compared across groups by two-sided Mann-Whitney U tests on the per-sample means, with Benjamini-Hochberg FDR correction applied across the three pairwise contrasts.

### SUPPLEMENTARY FIGURES

| Sample Name | Study Group | Sex | Age | Race | Ethnicity, Hispanic | Disease type | mRSS | Disease duration (years) | Medications | Sampled tissue, anatomical location | DV200 (%) |
|---|---|---|---|---|---|---|---|---|---|---|---|
| **HC01** | **Healthy** | F | 57 | White, Caucasian | No | None | n/a | n/a | None | Forearm | 84 |
| **HC02** | **Healthy** | F | 72 | White, Caucasian | No | None | n/a | n/a | None | Forearm | 73 |
| **HC03** | **Healthy** | F | 58 | White, Caucasian | Yes | None | n/a | n/a | None | Forearm | 89 |
| **HC04** | **Healthy** | F | 70 | White, Caucasian | No | None | n/a | n/a | None | Forearm | 69 |
| **HC05** | **Healthy** | F | 73 | White, Caucasian | No | None | n/a | n/a | None | Forearm | 74 |
| **HC06** | **Healthy** | F | 70 | White, Caucasian | No | None | n/a | n/a | None | Forearm | 65 |
| **SU01** | **Uninvolved** | F | 75 | White, Caucasian | Yes | lcSSc, CREST | 4 | 10 | None | Forearm | 57 |
| **SA01** | **Affected** | | | | | | | | | Finger, volar | 73 |
| **SU02** | **Uninvolved** | F | 42 | White, Caucasian | No | lcSSc, CREST | 8 | 12 | Plaquenil | Forearm | 83 |
| **SA02** | **Affected** | | | | | | | | | Forearm | 74 |
| **SU03** | **Uninvolved** | F | 71 | White, Caucasian | No | lcSSc, CREST | 4 | 16 | Mycophenolate | Forearm | 84 |
| **SA03** | **Affected** | | | | | | | | | Finger, unspecified | 78 |
| **SU04** | **Uninvolved** | F | 56 | Black, African American | No | lcSSc, CREST | 4 | 39 | Prednisone | Forearm | 80 |
| **SA04** | **Affected** | | | | | | | | | Finger, unspecified | 82 |
| **SU05** | **Uninvolved** | F | 77 | White, Caucasian | No | lcSSc, CREST | 8 | 21 | Cytoxan | Forearm | 76 |
| **SA05** | **Affected** | | | | | | | | | Finger, unspecified | 75 |
| **SU06** | **Uninvolved** | F | 73 | White, Caucasian | No | lcSSc, CREST | 2 | 1 | Trelegy, Sildenafil, Tadalafil, Levothyroxine | Forearm | 84 |
| **SA06** | **Affected** | | | | | | | | | Unspecified | 88 |
| **SU07** | **Uninvolved** | M | 66 | White, Caucasian | No | lcSSc, CREST | 4 | 4 | Omeprezole | Forearm | 83 |
| **SA07** | **Affected** | | | | | | | | | Finger, lateral | 88 |
| **SU08** | **Uninvolved** | F | 51 | White, Caucasian | Yes | lcSSc, CREST | 4 | 14 | Xeljanz, Tadalafil | Forearm | 88 |
| **SA08** | **Affected** | | | | | | | | | Finger, lateral | 89 |
| **SU09** | **Uninvolved** | F | 74 | White, Caucasian | No | lcSSc, CREST | 5 | 25 | Celebrex | Forearm | 77 |
| **SA09** | **Affected** | | | | | | | | | Finger, unspecified | 75 |

**Table S1. Cohort demographics, clinical characteristics, sampled tissue and quality metrics.** Full cohort table showing information collected at time of skin biopsy: Sample Name; Study Group; Sex; Age; Race; Ethnicity; Disease type; mRSS; Disease duration (years); Medications; Sampled tissue; DV200 (%). Healthy Control group, n=6, single skin biopsy; lcSSc group, n=9, two paired biopsies from clinically uninvolved and clinically affected skin tissue.

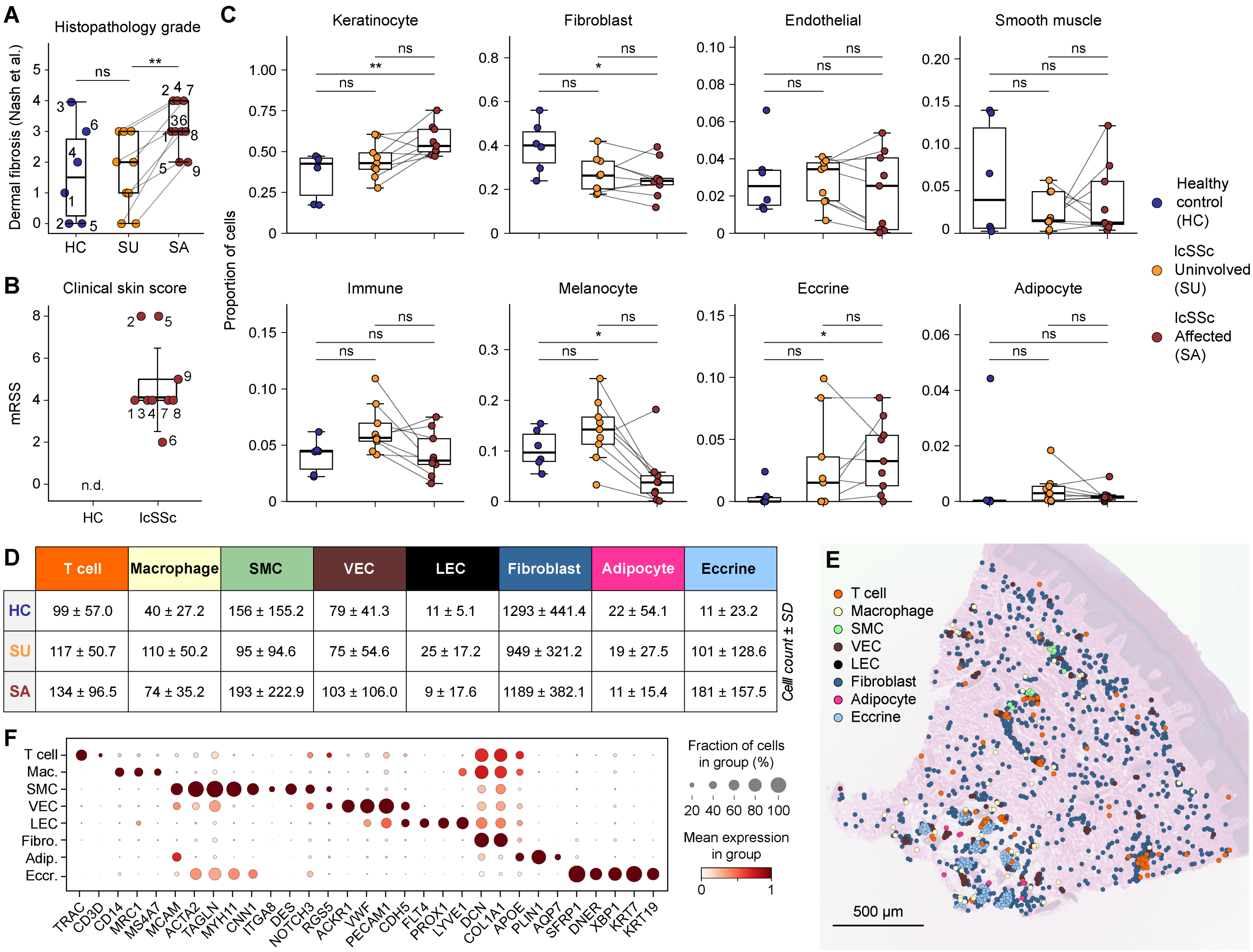


| | T cell | Macrophage | SMC | VEC | LEC | Fibroblast | Adipocyte | Eccrine |
|---|---|---|---|---|---|---|---|---|
| HC | 99 ± 57.0 | 40 ± 27.2 | 156 ± 155.2 | 79 ± 41.3 | 11 ± 5.1 | 1293 ± 441.4 | 22 ± 54.1 | 11 ± 23.2 |
| SU | 117 ± 50.7 | 110 ± 50.2 | 95 ± 94.6 | 75 ± 54.6 | 25 ± 17.2 | 949 ± 321.2 | 19 ± 27.5 | 101 ± 128.6 |
| SA | 134 ± 96.5 | 74 ± 35.2 | 193 ± 222.9 | 103 ± 106.0 | 9 ± 17.6 | 1189 ± 382.1 | 11 ± 15.4 | 181 ± 157.5 |

**Figure S1. Cohort histopathology and dermal cell populations.**
(**A**) Blinded dermal fibrosis grade by group according to an established scale (PMID 17452515)(0, no homogenization of collagen; 1, <25% sclerosis; 2, focal sclerosis, <50%; 3, 50-75% sclerotic change; 4, pan-dermal sclerosis; 5, pan-dermal sclerosis extending into the hypodermis with fibrous septa); numbers denote donors (HC vs SU, Mann-Whitney U, ns; SU vs SA, Wilcoxon signed-rank, p=.008; ordered trend by Jonckheere-Terpstra, p=.014). (**B**) Modified Rodnan skin score (mRSS) for lcSSc donors; n.d., not determined (HC). (**C**) Per-sample proportion of the eight broad cutaneous populations by group, paired SU-SA connected. (**D**) Mean ± SD cell counts per sample of finer immune, vascular, stromal, and exocrine populations by group. (**E**) Representative section with these finer populations overlaid on H&E image. (**F**) Dot plot of lineage markers for the populations in (D). Box plots, median and IQR; paired SU-SA connected; *p<.05, **p<.01. Scale bar, 500 µm.

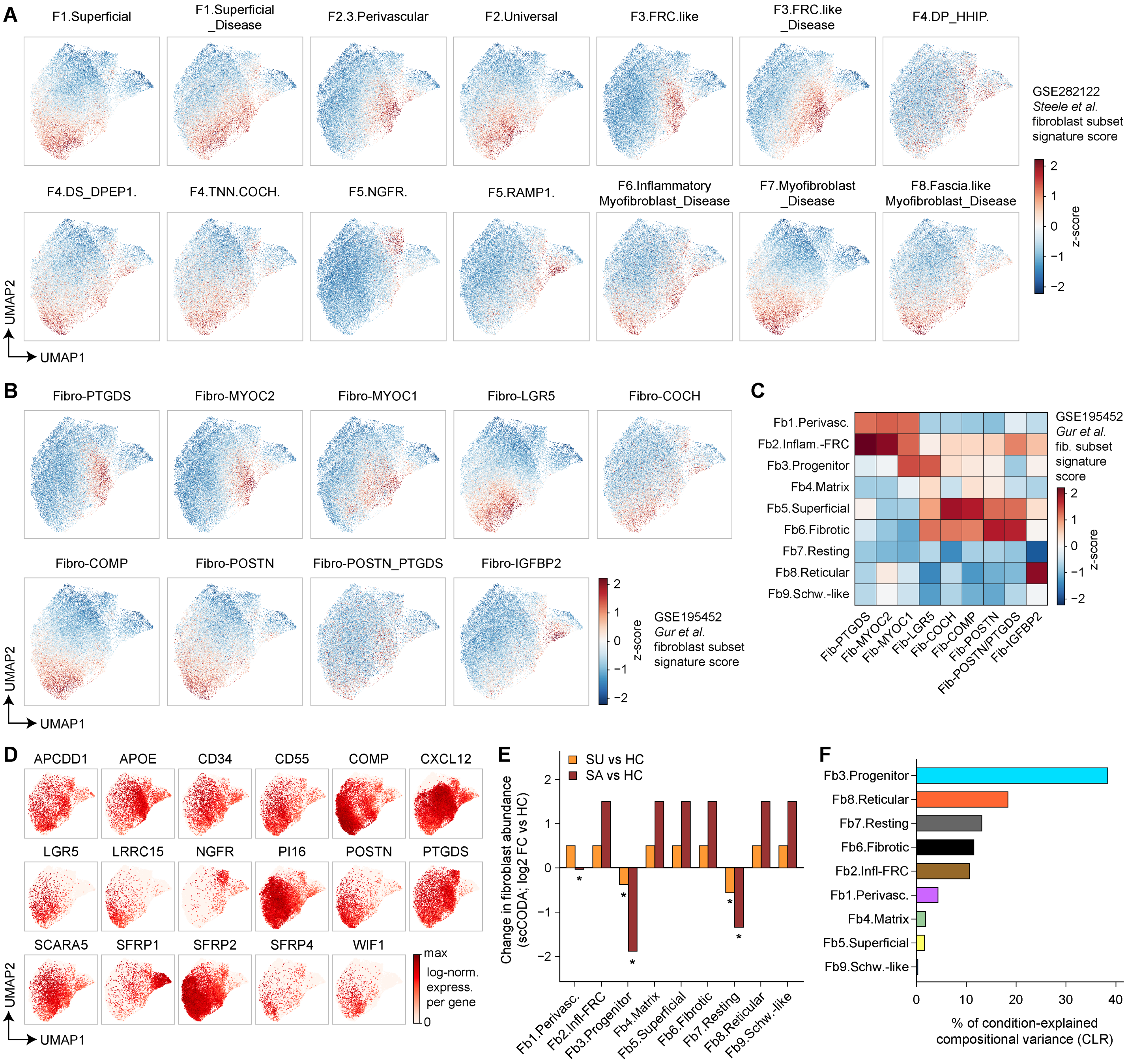


**Figure S2. The Fb3.Progenitor subset aligns with LGR5$^+$ fibroblasts perturbed in SSc.** (**A**) Steele et al. fibroblast signature scores projected onto the fibroblast UMAP. (**B**) Gur et al. fibroblast signature scores (9 populations) on the UMAP. (**C**) Enrichment (z-score) of Gur et al. populations across Fb1-Fb9. (**D**) Expression of lineage-defining and subset marker genes on the fibroblast UMAP. (**E**) scCODA change in fibroblast abundance (log2 fold change versus HC; SU and SA) per subpopulation; asterisks mark credible changes. (**F**) Percentage of condition-explained compositional variance (centered-log-ratio) contributed by each fibroblast subpopulation.

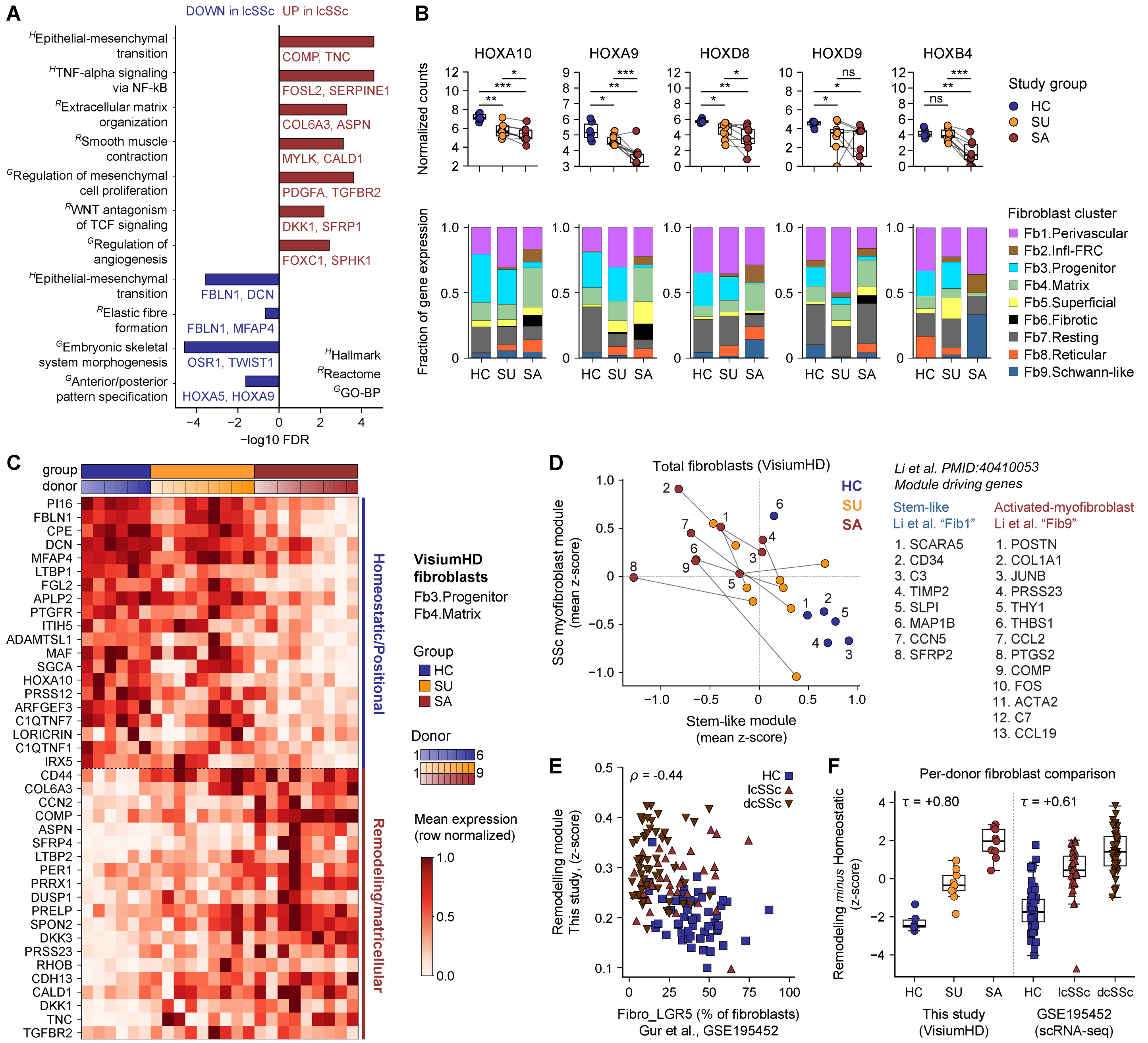


**Figure S3. Positional genes and stem-like programs are altered in clinically uninvolved skin.** (**A**) Pathway enrichment (-log10 FDR) of gene sets down- (blue) and up-regulated (red) in lcSSc fibroblasts (H, Hallmark; R, Reactome; G, GO-BP), with representative leading-edge genes. (**B**) Pseudo-bulk expression of homeobox genes (HOXA10, HOXA9, HOXD8, HOXD9, HOXB4) by group, with fibroblast-subpopulation contribution below. (**C**) Row-normalized expression of the 20 homeostatic/positional and 20 remodeling/matricellular module genes in Fb3.Progenitor and Fb4.Matrix across donors. (**D**) Per-sample Li et al. (PMID 40410053) stem-like (Fib1) versus activated-myofibroblast (Fib9) module scores, with module genes listed; numbers denote donors. (**E**) Matrix-remodeling module versus progenitor fraction (Fibro_LGR5) in Gur et al. (GSE195452), colored by group (Spearman ρ = −0.44). (**F**) Remodeling-minus-homeostatic module score by group in this study (HC, SU, SA) and in Gur et al. (HC, lcSSc, dcSSc) (within-cohort Kendall τ = +0.80 and +0.61). Box plots, median and IQR; *FDR<.05, **FDR<.01, ***FDR<.001 (B: limma-voom contrasts, Benjamini-Hochberg).

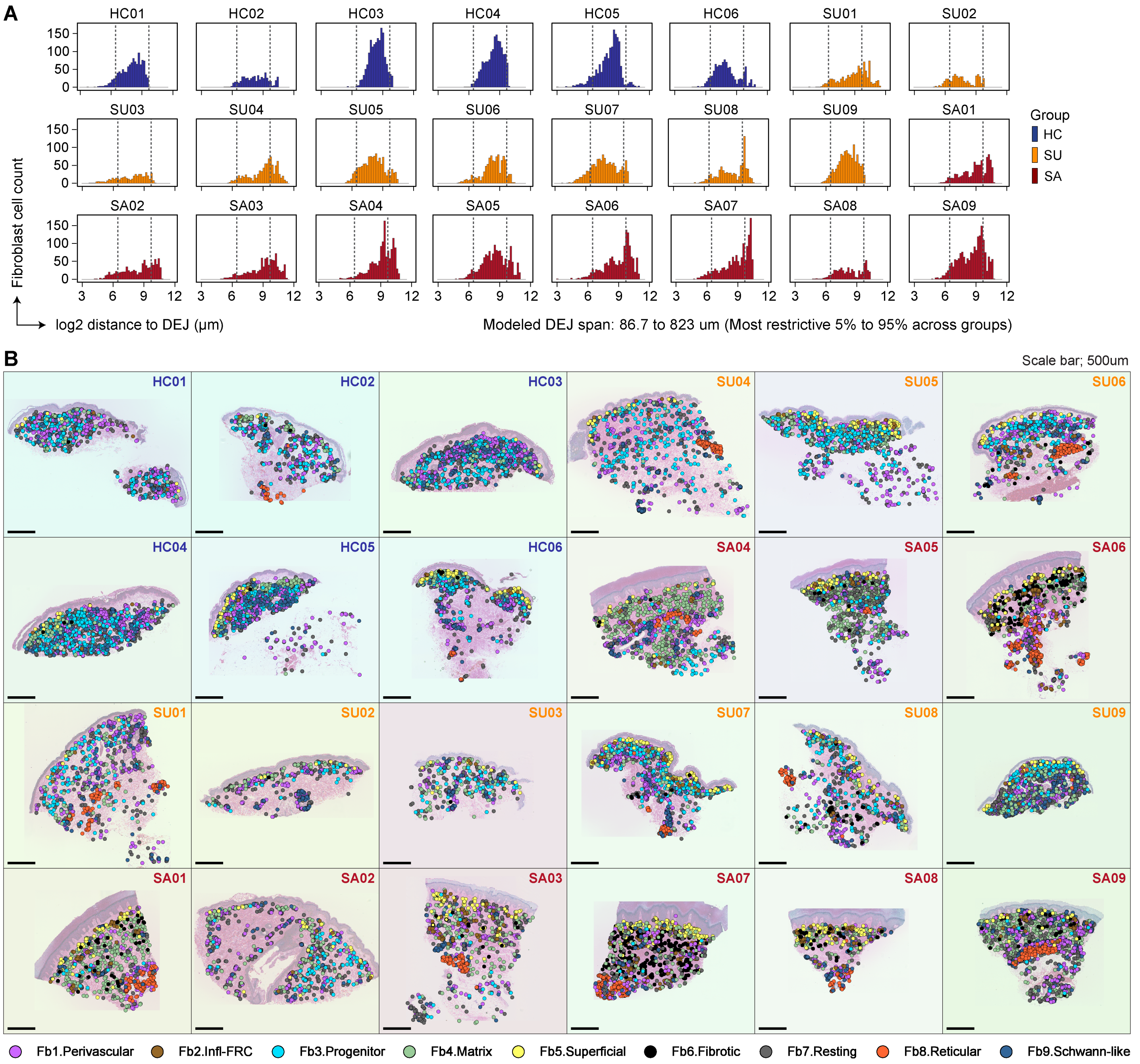


**Figure S4. Per-sample dermal fibroblast distributions and spatial maps.**
(**A**) Fibroblast count versus log2 distance from the DEJ for each of the 24 samples (dashed lines, 5th/95th percentiles; modeled DEJ span 86.7-823 µm, the most restrictive range across groups). (**B**) Spatial distribution of Fb1-Fb9 for all 24 samples, oriented epidermis-up. Scale bars, 500 µm.

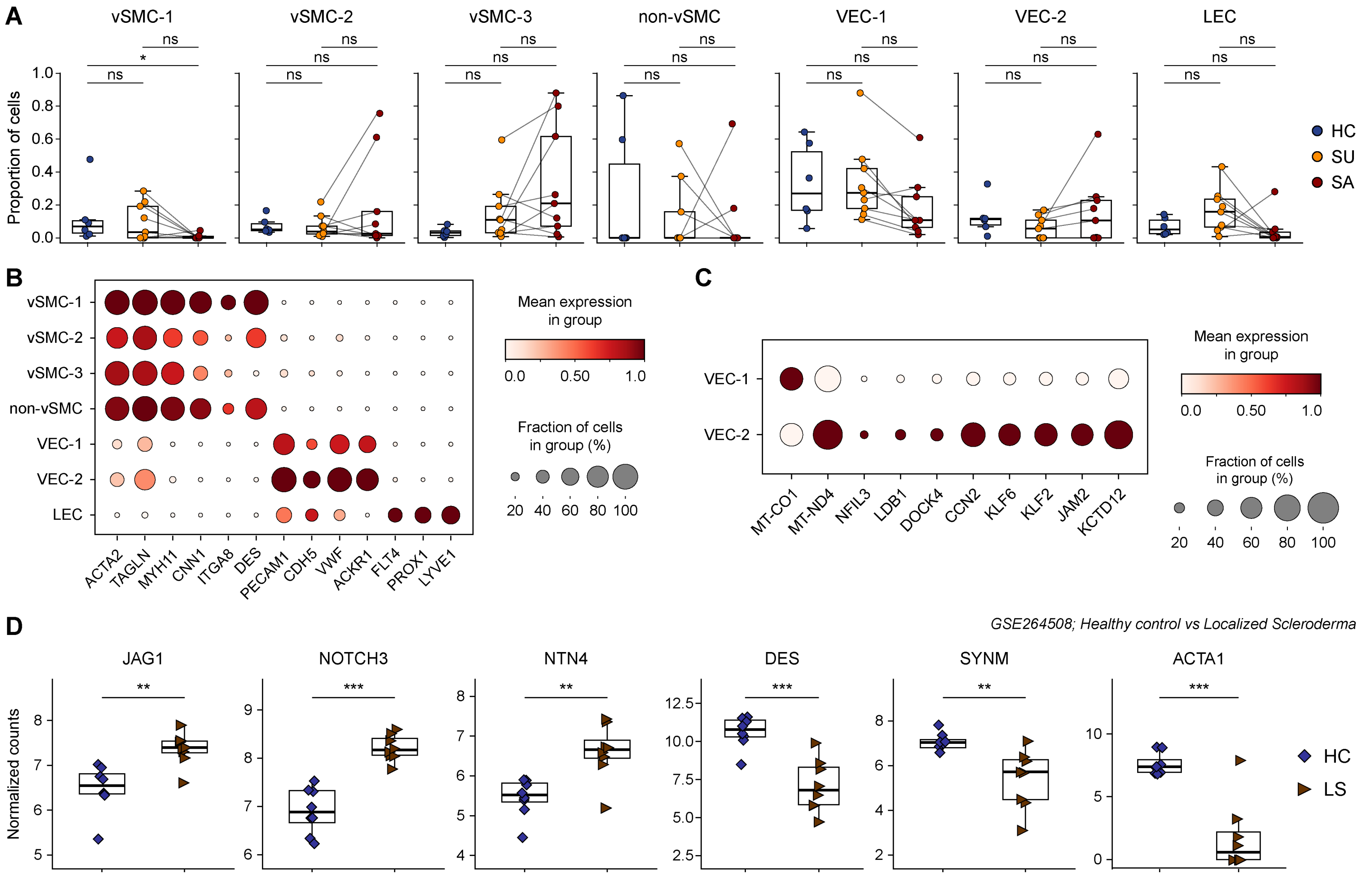


**Figure S5. Characterization of the dermal vascular smooth muscle and endothelial compartments.** (**A**) Per-donor proportion of each vascular subpopulation (vSMC-1/2/3, non-vSMC, VEC-1/2, LEC) by group, paired SU-SA connected. (**B**) Dot plot of vascular markers across the seven subpopulations: smooth-muscle (ACTA2, TAGLN, MYH11, CNN1, ITGA8, DES), vascular-endothelial (PECAM1, CDH5, VWF, ACKR1), and lymphatic-endothelial (FLT4, PROX1, LYVE1). (**C**) Dot plot of genes distinguishing VEC-1 from VEC-2. (**D**) External validation (GSE264508, healthy control, HC, versus localized scleroderma, LS): normalized counts of markers up-regulated in LS (JAG1, NOTCH3, NTN4) and down-regulated in LS (DES, SYNM, ACTA1). Box plots, median and IQR; HC comparisons, two-sided Mann-Whitney U test; SU vs SA, two-sided Wilcoxon signed-rank test on donor pairs; Benjamini-Hochberg correction; *p<.05, **p<.01, ***p<.001.

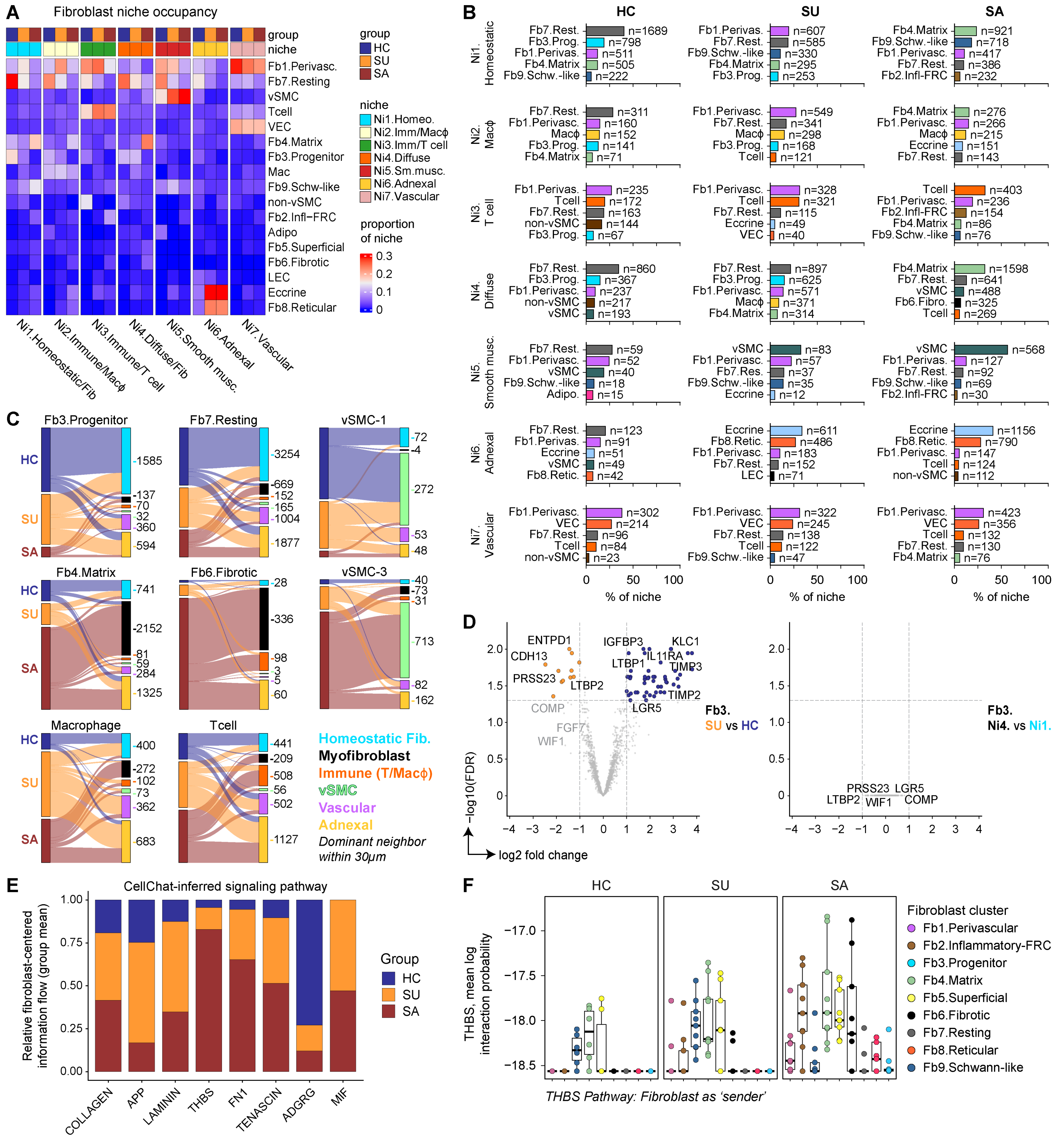


**Figure S6. Fibroblast neighborhood composition and inferred signaling changes in lcSSc.** (**A**) Niche occupancy (cellular composition) heat map by group. (**B**) Top 5 constituent populations of each niche per group (cell counts shown). (**C**) Alluvial plots of the dominant neighbor within 30 µm for representative focal populations across groups. (**D**) Volcano plots of Fb3.Progenitor differential expression by group (SU-vs-HC) and by niche (Ni4-vs-Ni1). (**E**) CellChat-inferred relative fibroblast-centered information flow (group mean) for the top signaling pathways (COLLAGEN, APP, LAMININ, THBS, FN1, TENASCIN, ADGRG, MIF). (**F**) THBS-pathway interaction probability with fibroblasts as sender, by subpopulation (Fb1-Fb9) and group.

**A**

| | SA02-Fb3$^{lo}$ vs SA$^{avg}$ | SA02-Fb3$^{hi}$ vs SU$^{avg}$ |
|---|---|---|
| Fb3.Progenitor (% fibs) | **1.9** \|\| 2.2 | **12.7** \|\| 12.7 |
| Fb4.Matrix (% fibs) | **14.9** \|\| 27.3 | **15.8** \|\| 9.2 |
| Fb2.Infl-FRC (% fibs) | **0.0** \|\| 6.2 | **1.4** \|\| 1.1 |
| Fb6.Fibrotic (% fibs) | **0.0** \|\| 5.0 | **0.0** \|\| 0.7 |
| vSMC-1 (% vasc) | **24.3** \|\| 2.5 | **16.2** \|\| 15.8 |
| vSMC-3 (% vasc) | **0.0** \|\| 40.7 | **1.2** \|\| 15.7 |
| Homeostatic module (z-score) | **-1.36** \|\| -0.54 | **-0.12** \|\| +0.09 |
| Remodeling module (z-score) | **-0.76** \|\| +0.82 | **-0.26** \|\| -0.17 |
| Li et al. stem-fib. (score) | **+7.89** \|\| +7.99 | **+8.73** \|\| +8.68 |
| Gur et al. SSc sig. (z-score) | **+0.56** \|\| +0.29 | **+0.03** \|\| -0.05 |
| Fb3.Prog. DEJ depth (µm) | **589** \|\| 757 | **553** \|\| 361 |
| Fb3.Prog. loneliness (µm) | **9.3** \|\| 9.6 | **10.2** \|\| 10.5 |
| Ni1.Homeostatic (% niches) | **14.9** \|\| 27.6 | **14.4** \|\| 23.1 |
| Ni4.Diffuse (% niches) | **33.8** \|\| 33.4 | **41.4** \|\| 33.6 |

**Figure S7. Adjacent regions within a clinically affected lcSSc sample resemble uninvolved and fibrotic skin.**

(**A**) Feature values for the two halves of SA02 (left of "||") compared with the matched group average (right of "||"): SA02-Fb3$^{lo}$ versus the SA average (left column) and SA02-Fb3$^{hi}$ versus the SU average (right column), covering fibroblast composition, transcriptional modules, Fb3.Progenitor position, niche occupancy, and vascular composition (SA average excludes SA02).
